# Multiple myeloma long-term survivors display sustained immune alterations decades after first line therapy

**DOI:** 10.1101/2023.05.27.542555

**Authors:** Raphael Lutz, Florian Grünschläger, Malte Simon, Marcus Bauer, Schayan Yousefian, Niklas Beumer, Lea Jopp-Saile, Mohamed H.S. Awwad, Georg Steinbuss, Anastasia Sedlmeier, Tobias Boch, Dominik Vonficht, Marc-Andrea Baertsch, Brian G.M. Durie, Niels Weinhold, Marc S. Raab, Claudia Wickenhauser, Andreas Trumpp, Carsten Müller-Tidow, Daniel Hübschmann, Benedikt Brors, Hartmut Goldschmidt, Charles D. Imbusch, Michael Hundemer, Simon Haas

**Author notes:** These authors contributed equally (Raphael Lutz, Florian Grünschläger, Malte Simon). shared correspondence. Conflict of interest: The authors declare no potential conflicts of interest. Corresponding authors: Correspondence should be addressed to Hartmut Goldschmidt, Charles D. Imbusch, Michael Hundemer or Simon Haas.

## Abstract

The long-term consequences of cancer or cancer therapy on the patients’ immune system years after cancer-free survival remain poorly understood. Here, we have performed an in-depth characterization of the bone marrow ecosystem of multiple myeloma long-term survivors at initial diagnosis and up to 17 years following cancer-free survival. Using comparative single-cell analyses in combination with molecular, genomic and functional approaches, we demonstrate that multiple myeloma long-term survivors display pronounced alterations in their bone marrow microenvironment associated with impaired immunity. These immunological alterations were frequently driven by an inflammatory immune circuit fueled by the long-term persistence or resurgence of residual myeloma cells. Notably, even in the complete absence of any detectable residual disease for decades, sustained changes in the immune system were observed, suggesting an irreversible ‘immunological scarring’ caused by the initial exposure to the cancer and therapy. Collectively, our study provides key insights into the molecular and cellular bone marrow ecosystem of multiple myeloma long-term survivors, revealing reversible and irreversible alterations of the immune compartment, which can serve as diagnostic and predictive tools.

**Statement of significance:** Large-scale single-cell profiling of a unique cohort of multiple myeloma long-term survivors uncovered that exposure to cancer and its treatment causes both reversible and irreversible immune alterations associated with impaired immunity. These findings have far-reaching implications for the understanding of long-term immune alterations in cancer, which need to be considered also in the context of immune therapeutic approaches. Furthermore, our study demonstrates how cancer-associated immune trafficking can be used to predict disease re-initiation in the bone marrow, opening new avenues for minimally invasive disease monitoring.

## Introduction

The immune system plays a key role in the prevention, development and treatment of cancer. Powerful immune surveillance mechanisms constantly monitor tissues to remove potentially cancerous cells. However, malignant tumors can evade immune control or even hijack immunological processes to propel tumor growth. Notably, the interaction between the tumor and the immune system induces bidirectional adaptations. Well studied examples for immunological changes induced by the continuous exposure to tumor cells include the exhaustion and dysfunction of T cells, as well as the suppressive polarization of myeloid immune cells, such as tumor associated macrophages or myeloid-derived suppressor cells [1-5]. In infectious diseases, irreversible immune dysfunction has been described, long after the infection has been cleared, a phenomenon termed immunological scarring [6, 7]. However, whether cancer or cancer treatment may cause similar long-term consequences on the immune system years after cancer-free survival remains poorly understood.

Multiple Myeloma (MM) is a hematologic neoplasm and is characterized by the clonal proliferation of malignant plasma cells within the bone marrow (BM). MM provides a prime example for a disease that depends on the interplay with its tumor microenvironment [8, 9]. Recent bulk and single-cell genomic efforts dissected the clonal complexity as well as clonal evolution patterns of MM from precursor stages to symptomatic disease and upon refractory cancer after multiple therapy lines [10-12]. While transcriptional stability has been observed in the transition from precursor states to MM progression, more dynamic shifts within the transcriptome and clonal outgrowth occurred upon refractory cancer [13]. Besides the genomic evolution of myeloma cells, substantial changes in the immune and stromal cell composition have been described across the different MM disease stages promoting an inflammatory BM microenvironment upon disease progression [8, 14]. Cell-cell interactions within the BM appear to be crucial to mediate tumor growth in MM highlighting the importance for a deeper understanding of the tumor ecosystem at different disease stages [15]. While recent studies on the MM ecosystem focused on disease progression from precursor stages as well as refractory disease, it remains unclear whether myeloma and myeloma therapy causes long-term alterations of the immune system years to decades after progression-free survival.

Despite improved therapy options, MM remains an incurable disease and only a minor fraction of MM patients experiences long-term survival (LTS) over 7 years after first line therapy [16, 17]. Nonetheless, even patients in complete remission (CR) without detectable measurable residual disease (MRD) may ultimately experience biochemical progression years after progression-free survival. Previous studies on the LTS phenomenon in MM focused on quantitative changes in immune cell types [18-20]. However, the transcriptional evolution patterns of myeloma cells in LTS patients as well as the long-term molecular adaptations of the BM microenvironment years after progression-free survival remain unexplored.

Here we have characterized the BM ecosystem of a unique patient group of MM long-term survivors at initial diagnosis (ID) and 7-17 years after first line therapy. Of note, LTS patients displayed sustained alterations in the immune microenvironment if compared to age-matched controls. These changes were associated with resurgence of disease activity but were also detectable in patients that were considered functionally cured, suggesting both reversible and irreversible long-term consequences of the disease and therapy. We identified bone marrow infiltrating inflammatory T cells as part of an inflammatory circuit, propelling these sustained immune aberrations. Importantly, this disease-associated immune cell trafficking can be used to reliably track the re-initiation of the disease.

## Results

### The bone marrow ecosystem of multiple myeloma long-term survivor patients

The long-term alterations of the immune system years to decades after a cancer diagnosis remain unknown. To elucidate the bone marrow ecosystem of LTS cancer patients, our study included 24 multiple myeloma patients who experienced LTS for 7 to 17 years (median 10.5 years) after first line therapy with standard induction regimen and high dose therapy followed by autologous stem cell transplantation (Fig.1a, Table 1). Notably, the favorable outcome of these patients could not have been predicted by state-of-the-art risk stratification tools, as 10 out of 24 patients displayed an intermediate, or poor prognosis according to the International Staging System (ISS) [21] and 4 patients even harbored high risk cytogenetic aberrations. Average myeloma cell infiltration within the BM across all patients at ID was remarkably high (mean 50%). For 11 of these MM patients with paired longitudinal samples at ID and upon LTS 7-17 years post diagnosis, we performed droplet-based single-cell RNA-sequencing (scRNAseq) of total BM mononuclear cells. In addition, CD3+ T cells were separately profiled in all cases by scRNAseq to ensure sufficient coverage of the T cell compartment, even in the presence of high tumor burden. Bone marrow samples from three healthy, age-matched donors were included as controls, applying the identical workflow (Fig.1a, Extended data Fig. 1a). Following data integration, clustering and dimensionality reduction across experiments, we analyzed 213,200 high-quality BM cells covering the vast majority of previously described hematopoietic cell types and cell states of the BM (Fig. 1b, Extended data Fig. 1b). These included plasma cells, all hematopoietic stem and progenitor cell stages, T cell and natural killer (NK) cell populations, several dendritic cell and monocyte subpopulations as well as the main B cell differentiation states.

**Figure 1.**
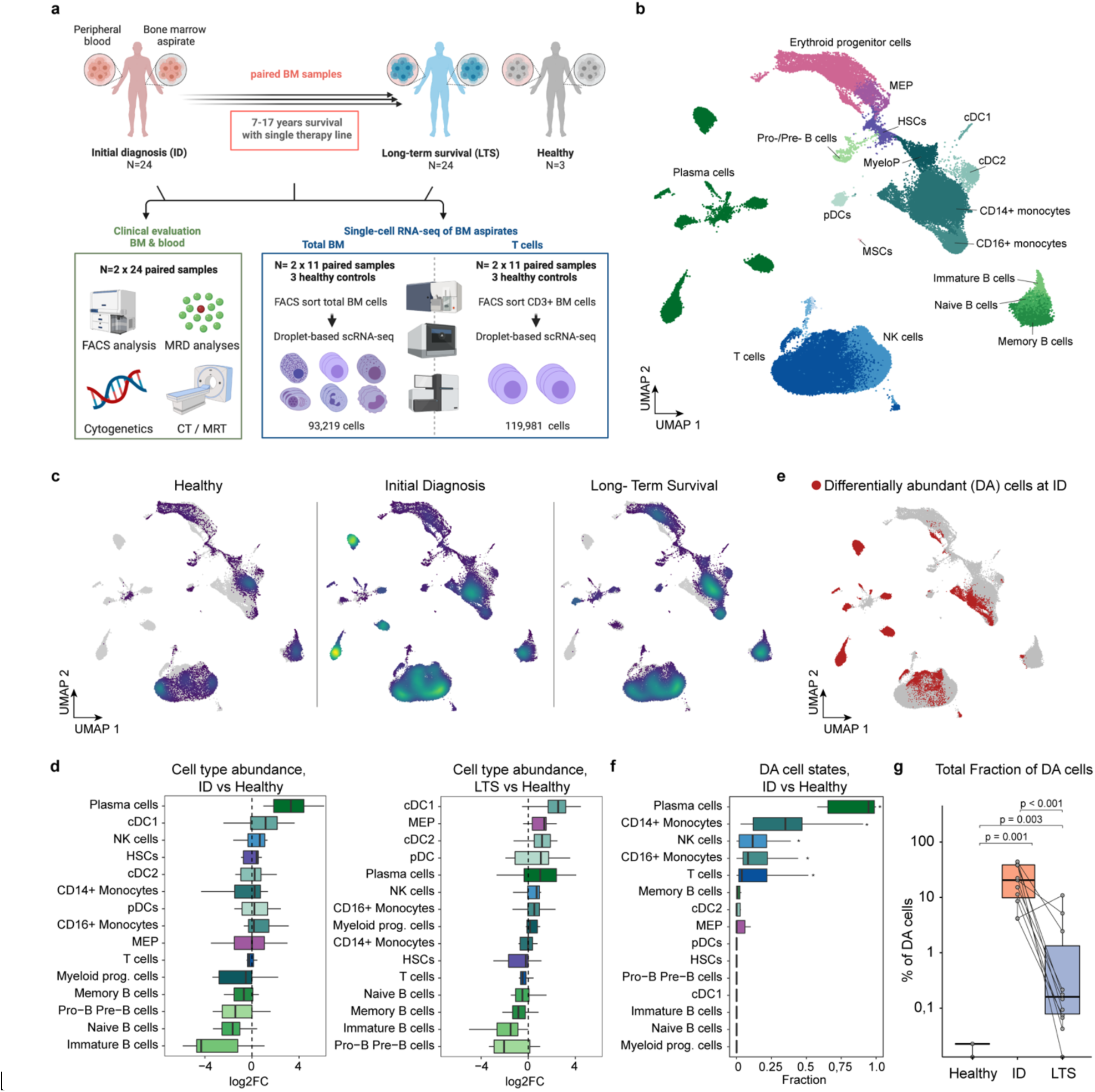
The bone marrow ecosystem of multiple myeloma long-term survivor patients. Also see extended data Figure 1. **(a)** Overview of the study design and experimental layout. **(b)** Global UMAP representation of scRNAseq data of paired human BM samples from 11 MM long-term survivor patients at initial diagnosis (ID) and long-term survival (LTS), as well as 3 healthy, age-matched controls. **(c)** Global UMAP split by clinical groups. The density and distribution of cells is color-coded. Grey represents all remaining cells. **(d)** Changes in cell type abundancies between ID or LTS in comparison to healthy donors **(e)** Global UMAP highlighting differentially abundant cells (red) determined by DA-Seq at initial diagnosis as compared to cells from healthy controls. **(f)** Fractions of differentially abundant cells (DA-cells) compared to all cells per cell type and patient at initial diagnosis. Benjamini-Hochberg (BH) adjusted significant differences (p < 0.05) evaluated by unpaired two-sided Wilcoxon rank sum test are highlighted. **(g)** Fractions of differentially abundant cells (DA-Seq) compared to all cells per patient within ID, LTS or healthy controls (Healthy). Dots represent sample means. BH corrected p-values from unpaired (Healthy/ID, Healthy/LTS) and paired (ID/LTS) two-sided Wilcoxon rank-sum tests are shown. Abbreviations: HSCs: hematopoietic stem cells, MEP: megakaryocyte-erythrocyte progenitors, MyeloP: myeloid progenitors, cDC1/2: conventional dendritic cells 1/2, pDCs: plasmacytoid dendritic cells, NK: natural killer cells, MSCs: mesenchymal stromal cells; ID: initial diagnosis, LTS: long-term survival. Box plots: center line, median; box limits, first and third quartile; whiskers, smallest/largest value no further than 1.5*IQR from corresponding hinge.

**Table 1:** Characteristics of patients with Multiple Myeloma in LTR: Patients highlighted in yellow were subjected to single cell RNA sequencing analysis, Abbreviations: PID: PatientID; ASCT= autologous stem cell transplantation; BM= bone marrow; CR= Complete remission; Mel = melphalan; NA = not available; n.a. not applicable; PC = plasma cells; PR= partial response; PAD= bortezomib – doxorubicin-dexamethasone; TAD = thalidomide-doxorubicin-dexamethasone; VAD= vincristine – doxorubicin – dexamethasone; VCD= bortezomib-cyclophosphamide-dexamethasone; VID= vincristine – ifosfamide – dexamethasone; VGPR= very good partial response

| Number | Patient ID | Gender (M/F) | Age | MM Type | CRAB criteria | Time after ASCT (years) | Stage (ISS) | Cytogenetics | Cytology: % PC in BM | Induction treatment | Pre-ASCT response | Conditioning | Maintenance, duration (in years after ASCT) | Post-ASCT response | Relapse from CR in 2018 (years after ASCT) | Sustained CR til 2022 |
| --- | --- | --- | --- | --- | --- | --- | --- | --- | --- | --- | --- | --- | --- | --- | --- | --- |
| 1 | P018 | M | 68 | IgG kappa | bone disease | 14 | I | standard | 5% | 3x VAD | NA | 2x Mel 200 | interferon, 8 | CR | 7 | No |
| 2 | P010 | M | 73 | IgG lambda | bone disease, anemia | 11 | I | standard | 30% | 3x VAD | PR | 2x Mel 200 | thalidomide, 2 | CR | 3 | No |
| 3 | P020 | F | 71 | IgA/IgG lambda | bone disease, anemia | 10 | III | standard | 90% | 3x VAD | VGPR | 2x Mel 200 | thalidomide, 1 | CR | 8 | No |
| 4 | P001 | F | 69 | IgG kappa | bone disease | 9.5 | I | standard | 15% | 3x VAD | PR | 1x Mel 200 | thalidomide, 2 | CR | 9 | No |
| 5 | P021 | F | 73 | IgG kappa | bone disease | 9 | II | high risk (del17p) | 50% | 3x PAD | VGPR | 2x Mel 200 | bortezomib, 2 | CR | 7 | No |
| 6 | P017 | M | 77 | IgG kappa | bone disease | 10 | I | standard | 10% | 3x VAD | PR | 1x Mel 200 | thalidomide, 1 | CR | 10 | No |
| 7 | P013 | M | 73 | IgG kappa | anemia | 9 | I | standard | 20% | 3x PAD | VGPR | 2x Mel 200 | none | CR | 6 | No |
| 8 | P009 | F | 56 | IgA lambda | bone disease | 9 | I | high risk (del 17p) | 60% | 3x PAD | VGPR | 2x Mel 200 | bortezomib, 2 | CR | 6 | No |
| 9 | P015 | F | 67 | IgA kappa | anemia | 9 | I | standard | 80% | 3x TAD | VGPR | 1x Mel 200 | None | VGPR | n.a. | No |
| 10 | P025 | M | 54 | BJ kappa | bone disease | 15 | NA | NA | NA | 3x VAD | NA | 2x Mel 200 | interferon, 2 | CR | n.a. | No |
| 11 | P022 | F | 58 | IgG kappa | bone disease | 14 | II | NA | 60% | 3x TAD | NA | 2x Mel 200 | thalidomide, 4 | CR | n.a. | No |
| 12 | P003 | M | 69 | IgG kappa | bone disease | 14 | III | standard | 80% | 3x TAD | PR | 2x Mel 200 | thalidomide, 4 | CR | n.a. | Yes |
| 13 | P004 | M | 70 | IgA lambda | bone disease | 9 | II | high risk (del 17p) | 100% | 3x PAD | nCR | 2x Mel 200 | bortezomib, 2 | CR | n.a. | No |
| 14 | P016 | F | 65 | BJ kappa | bone disease | 11 | I | standard | 20% | 3x PAD | VGPR | 2x Mel 200 | bortezomib, 2 | CR | n.a. | Yes |
| 15 | P023 | F | 79 | IgA lambda | bone disease | 14 | I | standard | 30% | 3x VAD | CR | 2x Mel 200 | interferon, NA | CR | n.a. | Yes |
| 16 | P011 | F | 58 | IgG kappa | bone disease | 17 | NA | NA | 80% | 4x VID | NA | 2x Mel 200 | interferon, 13 | CR | n.a. | Yes |
| 17 | P005 | M | 59 | IgG kappa | renal failure, anemia | 12 | II | standard | 20% | 3x VAD | PR | 2x Mel 200 | interferon, NA | CR | n.a. | Yes |
| 18 | P002 | M | 65 | IgG lambda | bone disease | 11 | I | standard | 70% | 3x VAD | VGPR | 2x Mel 200 | thalidomide, 2 | CR | n.a. | No |
| 19 | P007 | M | 75 | IgA lambda | bone disease | 11 | I | standard | 80% | 3x PAD | VGPR | 2x Mel 200 | bortezomib, 2 | CR | n.a. | Yes |
| 20 | P012 | F | 61 | IgG kappa | bone disease | 11 | III | standard | 30% | 3x VAD | PR | 1x Mel 200 | thalidomide, 3 | CR | n.a. | Yes |
| 21 | P006 | M | 55 | IgG kappa | bone disease | 10 | II | high risk (gain 1q21) | 50% | 3x PAD | VGPR | 2x Mel 200 | bortezomib, 2 | CR | n.a. | Yes |
| 22 | P024 | M | 68 | IgG lambda | renal failure | 9.5 | III | high risk (t4;14) | 80% | 3x PAD | VGPR | 2x Mel 200 | bortezomib, 2 | CR | n.a. | No |
| 23 | P014 | M | 60 | IgG kappa | bone disease, hypercalcemia | 9 | I | standard | 30% | 3x TAD | CR | 1x Mel 200 | None | CR | n.a. | Yes |
| 24 | P026 | F | 46 | IgG lambda | bone disease | 7 | II | standard | 50% | 3x VCD | PR | 2x Mel 200 | lenalidomide, 1 | CR | n.a. | No |

Comparing immune cell compositions of healthy donors with patients at ID revealed an expected enrichment for plasma cells and a trend towards higher amounts of cDC1 and NK cells, as well as a depletion of different B cell stages as described by previous studies (Fig. 1c,d, Extended data Fig. 1c,d) [10, 22]. At the LTS timepoint, the BM composition was partially normalized, however a significant enrichment of the dendritic cell compartments cDC1 and cDC2 constituted a unique feature of LTS patients (Extended data Fig. 1d). Beside changes in the BM cell type composition, we also observed considerable transcriptional perturbations within many BM-resident cell types, reflecting disease-associated adaptations of cellular transcriptomic states (Fig.1c). To quantify these changes in cellular states associated with ID and LTS, we made use of DA-seq, a computational tool that measures how much a cell’s neighborhood is dominated by a certain biological state (see methods). As expected, a major transcriptomic remodeling from healthy to malignant plasma cells was observed at ID (Fig.1e,f). In addition, significant transcriptomic changes occurred within CD14+ monocytes, CD16+ monocytes as well as T and NK cells. Importantly, while the transcriptomic remodeling of immune cells partially normalized during LTS, which was in line with a reduced cancer cell burden in the BM, sustained signs of immune remodeling were maintained even decades after a single therapy line (Fig. 1g).

### Malignant plasma cells frequently persist during long-term survival and display a transcriptionally stable phenotype

Recent studies reported dynamic transcriptional shifts of malignant plasma cells and clonal outgrowth during disease courses induced by therapeutic interventions [13]. However, it remains poorly understood whether plasma cells driving relapse years after tumor-free survival undergo molecular adaptations in the absence of any therapy pressure. Moreover, it is unclear whether malignant plasma cells persist in the BM of LTS patients that are considered functionally cured.

To address these questions, we performed an in-depth analysis of plasma cells to explore the longitudinal changes of the tumor cell compartment throughout LTS. The transcriptional heterogeneity of the plasma cell compartment was reflected by patient-specific MM cell clusters and a cluster of putative healthy plasma cells to which all patients and the healthy controls contributed (Fig. 2a). Patient-specific clusters showed distinct gene expression patterns in line with published bulk RNA gene expression signatures, highlighting the diversity of our patient cohort (Extended data Fig. 2.1a) [23]. As expected, the expanded plasma cell compartment at ID partially normalized upon LTS. However, some patients still harbored a high fraction of plasma cells at the LTS state (Fig. 2b). To delineate healthy and malignant plasma cells, we analyzed copy number aberrations (CNA) using inferCNV (see methods, Extended data Fig. 2.2). Overall, 59 out of 63 CNAs detected by cytogenetics could also be identified by our single-cell analyses, permitting a clear discrimination between healthy and malignant plasma cells (Fig. 2c, Extended data Fig. 2.1b,c). Furthermore, plasma cells classified as malignant almost exclusively expressed a single immunoglobulin light chain, whereas plasma cells classified as healthy contained both kappa and lambda expressing cells, confirming the accuracy of our CNA analyses (Fig. 2d,e, Extended data Fig. 2.1e). The fraction of malignant plasma cells within the overall plasma cell pool (termed ‘malignancy score’) was increased in LTS patients that had experienced a biochemical progression from complete remission (CR) after a long-term remission phase, hereafter termed non-CR patients (Fig. 2f, Extended data Fig. 2.1d). As expected, patients that were in clinical CR harbored less or no malignant plasma cells. Moreover, the fraction of malignant cells defined by CNAs correlated with the result obtained from next generation flow cytometry for detection of measurable residual disease (MRD) (Fig. 2g). Nonetheless, our scRNAseq-based analysis was able to detect malignant plasma cells in patients previously classified as ‘Flow MRD negative’ at the LTS state highlighting the potential of single-cell genomics to detect and characterize such rare residual malignant cells.

**Figure 2.**
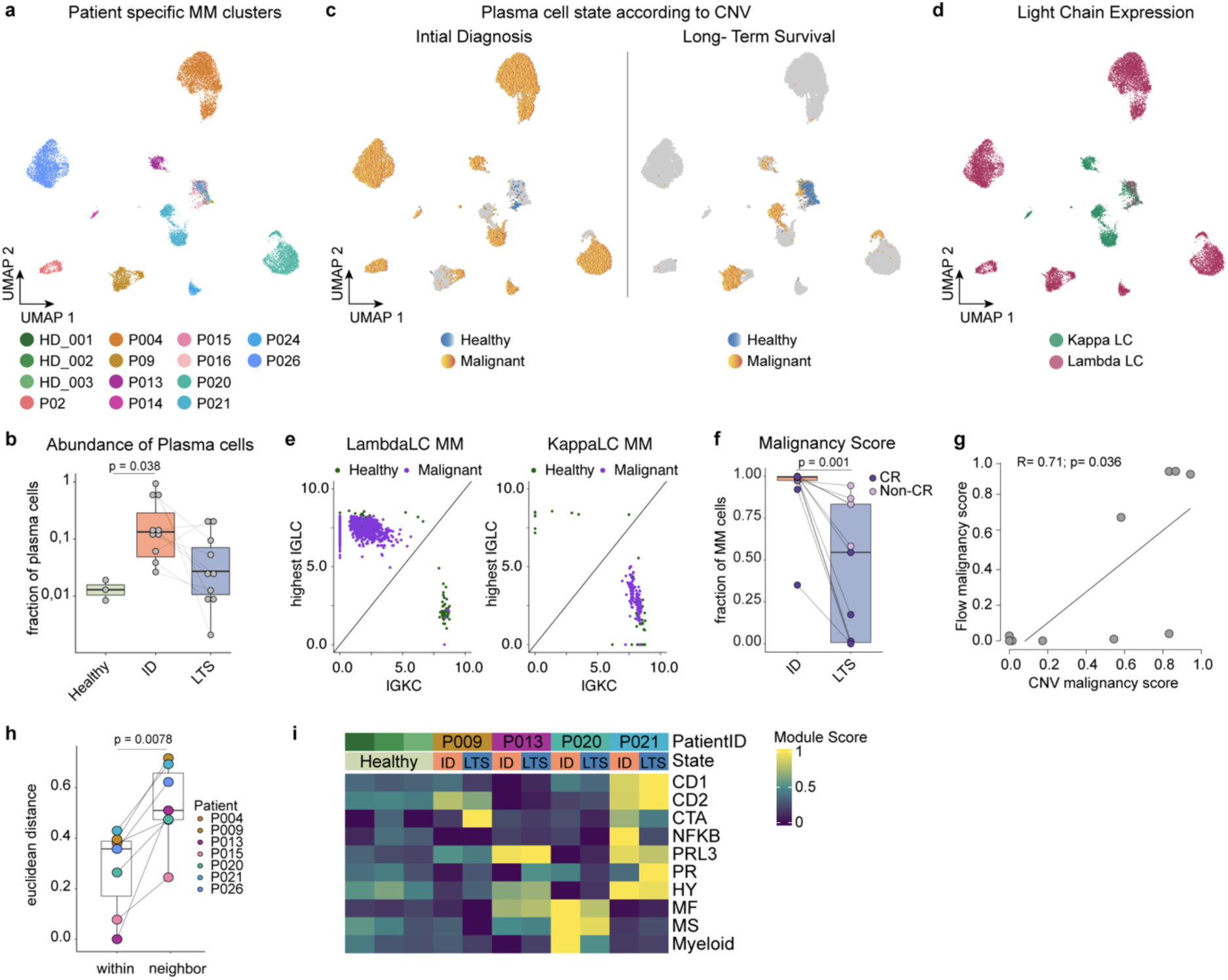
**Malignant plasma cells frequently persist during long-term survival and display a stable transcriptional phenotype.** Also see extended data Figures 2.1 and 2.2. **(a)** UMAP embedding of BM plasma cell (PC) compartment colored by patient. **(b)** PC fraction of total BM cells summarized by patient and compared between clinical groups (Healthy, ID, LTS). Dots indicate PC fraction of total BM cells for each sample. Significance was tested by unpaired Wilcoxon rank sum test. **(c)** Split UMAP of PCs by clinical groups (ID, LTS) highlighting their malignancy annotation (healthy, malignant) derived from inferCNV. Remaining cells from the respective other state are grayed out. **(d)** PC UMAP highlighting the dominant immunoglobulin light chain expression (Kappa: green; Lambda: red). **(e)** Representative scatter plots showcasing the immunoglobulin expression (highest lambda chain (IGLC) and kappa chain (IGKC)) of healthy (green) and malignant (violet) PCs. **(f)** Malignancy score (malignant PC fraction of total PCs) per patient at ID and LTS. Large dots indicate malignant PC fraction of total BM cells for each sample. Significance was tested by paired Wilcoxon signed rank test. **(g)** Correlation of malignancy score derived from Next Generation Flow MRD (number of Light Chain restricted plasma cells/all plasma cells) on y-axis with malignancy score derived from inferCNV analysis (number of malignant cells/all plasma cells) on x-axis. Spearman’s Rho and the significance level of correlation are indicated. **(h)** Euclidean distance of malignant plasma cells between ID and LTS within the same patient compared to the Euclidean distance of malignant plasma cells at ID and the respective nearest neighbor within top 30 principal components. Dots indicate the average Euclidean distance of each sample. Patients with less than 2 cells within one of the clinical states were excluded. Significance was tested by paired Wilcoxon signed rank test. **(i)** Heatmap showing average expression patterns (module scores; scaled by row) of known bulk RNA signatures [23] per patient and clinical state. Abbreviations: PC: plasma cells; ID: initial diagnosis; LTS: long-term survival; IGLC: immunoglobulin light chain; LC: lambda chain; KC: kappa chain. Box plots: center line, median; box limits, first and third quartile; whiskers, smallest/largest value no further than 1.5*IQR from corresponding hinge.

The mapping of CNAs in the single-cell data of the plasma cell compartment enabled us to address the question of how myeloma cells develop throughout the LTS state upon recurring disease activity. Malignant myeloma cells from the same patient at ID and LTS shared the highest transcriptional similarity to each other in comparison to myeloma cells from other patients (Fig. 2c,h). This suggested a high transcriptional stability of plasma cells upon resurgence of disease activity even after long lasting remission over years to decades. However, minor adaptations in the transcriptomic makeup between matched malignant plasma cells at ID and LTS were observed, as indicated by minor, but specific changes in the UMAP representation (Fig. 2c). To further study the molecular adaptations of myeloma cells, we focused on 4 patients with sufficient malignant cells captured for both matching clinical states to reliably obtain the subclonal composition of the respective patients (Extended data Fig. 2.2). Notably, we observed a changing subclonal composition which translated into specific changes of gene expression pattern of published transcriptomic signatures that are commonly used to categorize transcriptional patterns of myeloma cells (Fig. 2i) [23]. For example, P009 gained a cancer testis antigen (CTA) expression pattern, which is reported to be associated with a proliferative myeloma disease, whereas P021 lost the previously expressed NFKB signature upon resurgence of disease (Fig.2i). Together, our observations demonstrate that malignant plasma cells frequently persist in LTS patients and display an overall transcriptionally stable phenotype that is maintained for decades, while specific transcriptomic adaptions may occur.

### Multiple myeloma long-term survivor patients display sustained signs of immune remodeling decades after a single therapy line

While specific compositional changes in the BM microenvironment of LTS patients have been reported, it remains unknown whether these cell types adopt a cellular state similar to healthy BM cells or maintain signs of their current or past exposure to malignant plasma cells or therapy. Our initial analyses revealed a major transcriptomic remodeling of BM-resident immune cells during the disease course, with monocytic, T and NK cell compartments displaying the most extensive alterations in cell states besides the plasma cell compartment (Fig.1f). To further investigate these molecular changes across the clinical states, we first focused on the most remodeled cell compartment, classical CD14+ monocytic cells (Fig 3a). In line with our global DA-seq analysis, the majority of monocytes from ID patients clustered separately from monocytes of healthy donors, reflecting a disease-associated transcriptomic remodeling. Notably, this remodeling partially normalized in the LTS state, although a considerable number of monocytes maintained a remodeled state years to decades after a single, successful therapy line (Fig. 3a). To quantify the transcriptionally perturbed cells in the diseased states, we introduced a ‘dissimilarity score’ measuring whether a cell’s neighborhood is dominated either by the healthy or the disease state. Combining the dissimilarity score with machine learning-based approaches enabled us to classify cells as ‘healthy-like’ or ‘aberrant-like’ with high accuracy and a low false prediction rate (see methods). These analyses revealed that classical monocytes from patients at ID showed a high degree of dissimilarity to healthy monocytes and were frequently classified as ‘aberrant-like’. Upon LTS, only a partial normalization was observed, revealing a sustained transcriptional remodeling throughout LTS in a subset of monocytes (Fig. 3b-c). Of note, this remodeling pattern was associated with plausible biological processes as demonstrated in the next section.

**Figure 3.**
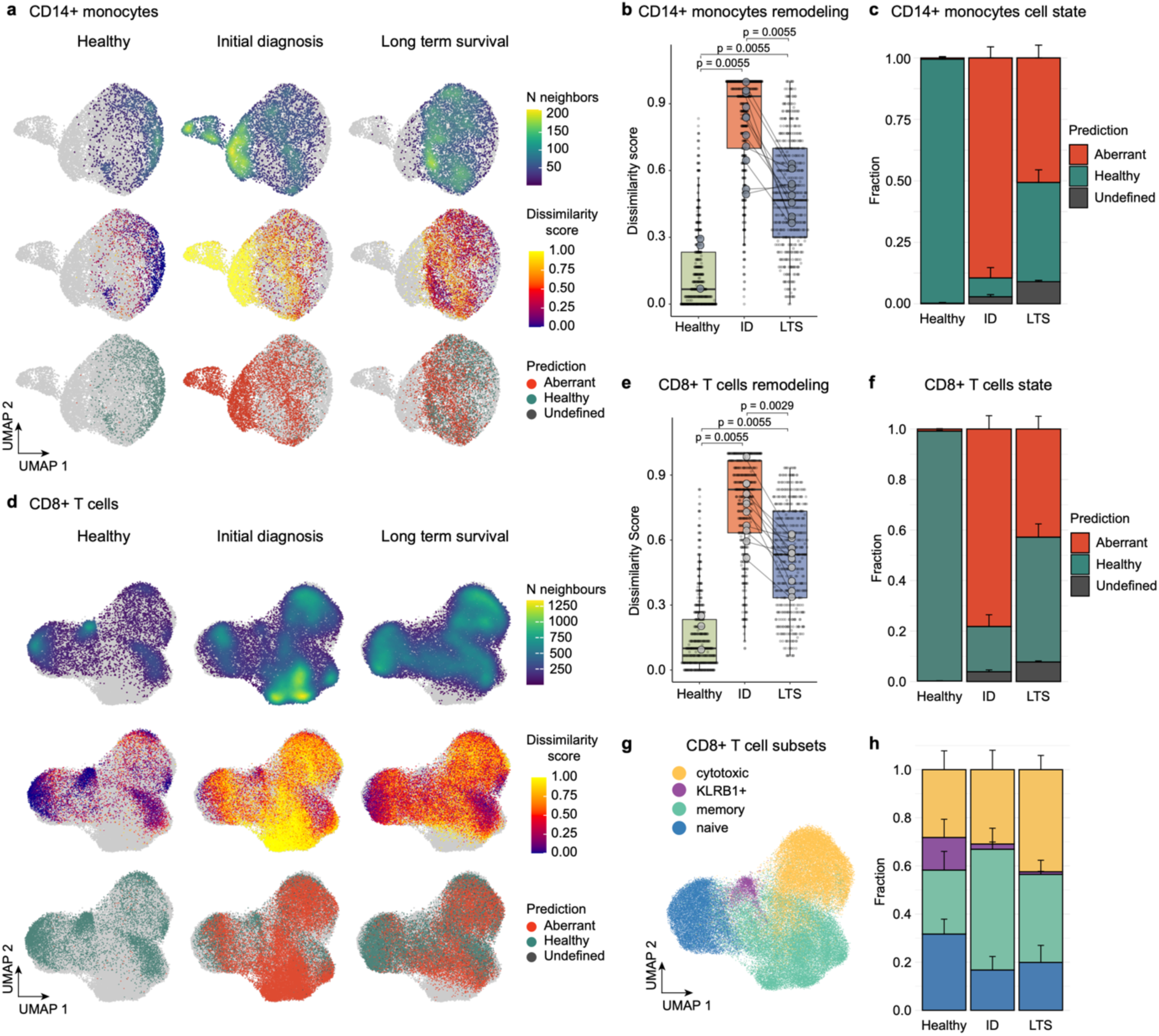
**Multiple myeloma long-term survivor patients display sustained signs of immune remodeling decades after a single therapy line.** Also see extended data Figure 3. **a)** UMAP of CD14+ monocytes from the BM dataset split by clinical groups. Cells are colored by the density (top row), dissimilarity score (middle row) and dissimilarity-based classification into aberrant-like, healthy-like and undefined cell states (bottom row). Cells from the respective other clinical states are depicted in grey. **(b)** Distribution of the dissimilarity score by clinical group summarizing the remodeling of CD14+ monocytes. Large dots indicate sample means. Benjamini-Hochberg adjusted p-values from unpaired (Healthy/ID, Healthy/LTS) and paired (ID/LTS) two-sided Wilcoxon rank-sum tests are shown. **(c)** Bar plot summarizing fractions of predicted cell states by clinical group from a. **(d)** UMAP of CD8+ T cells split by clinical groups. Cells are colored by the density (top row), dissimilarity score (middle row) and dissimilarity-based classification into aberrant-like, healthy-like and undefined cell states (bottom row). Cells from the corresponding other clinical states are shown in a grayscale. **(e)** Distribution of the dissimilarity score by clinical group summarizing the remodeling of CD8+ T cells. Large dots indicate sample means. Benjamini-Hochberg adjusted p-values from unpaired (Healthy/ID, Healthy/LTS) and paired (ID/LTS) two-sided Wilcoxon rank-sum tests are shown. **(f)** Bar plot summarizing fractions of predicted cell states by clinical group from d. **(g)** UMAP of CD8+ T cells, classified into naïve, memory, cytotoxic and KLRB1+ subsets. **(h)** Bar plot summarizing fractions of cell subsets by clinical group from g. Abbreviations: BM: bone marrow; ID: initial diagnosis; LTS: long-term survival. Bar plots: Error bars indicate the standard error of the mean (SEM); Box plots: center line, median; box limits, first and third quartile; whiskers, smallest/largest value no further than 1.5*IQR from corresponding hinge.

To investigate whether also other immune cell types display sustained transcriptional changes in the LTS state, we next focused on the T cell compartment. CD8+ T cell states were annotated in naive, memory, effector as well as KLRB1+ cells based on known transcriptomic marker genes (Fig. 3g, Extended data Fig.3a-c). Notably, also in the CD8+ T cell compartment a sustained transcriptional remodeling was observed upon long-term survival (Fig. 3d-f). Moreover, a significant and irreversible depletion of KLRB1+ CD8+ T cells was observed at the ID state and maintained throughout LTS (Fig. 3h).

In line with our observations from the classical monocyte and CD8+ T cell compartments, we observed a remodeling of non-classical CD16+ monocytes, as well as the CD4+ T and NK cell states at ID, which was partially sustained throughout LTS (Extended data Fig. 3d-o). Together, our data reveals a major remodeling of cell states across the majority of bone marrow cell types during active MM disease, which is sustained in a subset of cells throughout long-term survival.

### An inflammatory circuit underlies immune remodeling during active disease and long-term survival

To characterize disease-associated molecular programs responsible for the acute remodeling in the bone marrow ecosystem at ID, we performed a comprehensive gene set enrichment analysis (GSEA) comparing aberrant-like cell states with cells from healthy controls within all cell types of the bone marrow that displayed disease-associated remodeling. This analysis revealed a globally up-regulated inflammatory program (Hallmark TNFA signaling via NFKB and Hallmark inflammatory response) shared across all remodeled BM cell types, as well as cell type-specific changes (Fig. 4a). In particular, aberrant monocytes acquired a pro-inflammatory phenotype. The expression of inflammatory genes in monocytes correlated with their dissimilarity to healthy monocytes, peaked in ID patients and partially reversed throughout LTS (Fig. 4b). However, the remaining ‘aberrant-like’ monocytes in the LTS state specifically displayed a sustained inflammatory phenotype, suggesting a persistent inflammatory response of the classical monocyte compartment even decades after the first line therapy (Fig. 4c). As part of the inflammatory response, ‘aberrant-like’ monocytes displayed an increased chemokine activity and produced increased levels of proinflammatory cytokines and chemokines, including *CCL3*, *IL1B* and *CXCL8*, with the latter two known to support myeloma cell growth and survival (Fig. 4d-e, Extended data Fig. 4a-e) [24]. Interestingly, the corresponding receptors of *CXCL8, CXCR1* and *CXCR2* were mainly expressed on NK cells suggesting a role for *CXCL8* in the regulation and induction of leukocyte migration as reported previously (Extended data Fig. 4f) [25]. NK cells themselves switched from a cytotoxic to an inflammatory phenotype with increased chemokine activity, which was maintained throughout the LTS state (Fig. 4a,f).

To explore the interaction network between plasma cells and their microenvironmental cells at ID, we used CellPhoneDB to infer intercellular communications (see methods). We observed the highest number of interactions between myeloid and plasma cells (Fig. 4g). Notably, these interactions were significantly increased between remodeled CD14+ monocytes and plasma cells, suggesting that the remodeled state of CD14+ monocytes may be mediated by the interaction with plasma cells (Extended data Fig. 4g).

Importantly, remodeled T and NK cells were the main producers of the proinflammatory master cytokine interferon-gamma (*IFNg*) both at ID and LTS (Fig. 4h-I, Extended data Fig. 4m). Moreover, remodeled T and NK cells displayed significantly increased expression of the inflammatory chemokines *CCL3, CCL4* and *CCL5*, suggesting that they act as major regulators of the acute and sustained BM inflammation (Extended data Fig. 4h-j). In line with an increased synthesis of proinflammatory cytokines, including *IFNg*, by aberrant lymphocytes, we observed the strongest *IFNg* response in aberrant myeloid cells, including CD14+ and CD16+ monocytes as well as cDC2s (Fig. 4a). Notably, the IFN-inducible chemokines *CXCL9, CXCL10* and *CXCL11* were mainly expressed by CD16+ monocytes peaking at ID and being maintained at lower level throughout LTS (Extended data Fig. 4k,l). Aberrant *IFNg* expressing CD8+ T cells and NK cells specifically expressed *CXCR3*, the chemokine receptor mediating migration towards *CXCL9/10/11* sources, which we will elucidate in detail in the next section (Fig. 4j, Extended data Fig. 4n).

In summary, these data suggest that upon MM disease activity in the BM, inflammatory signals drive a positive feedback loop with IFNg secretion by aberrant lymphocytes inducing the release of *CXCL9/10/11* from myeloid cells (Extended data Fig. 5a). This in turn may lead to the recruitment of *CXCR3+* inflammatory CD8+ T cells to the BM (see below) causing an inflammatory circuit which is maintained at a lower level in LTS patients.

**Figure 4.**
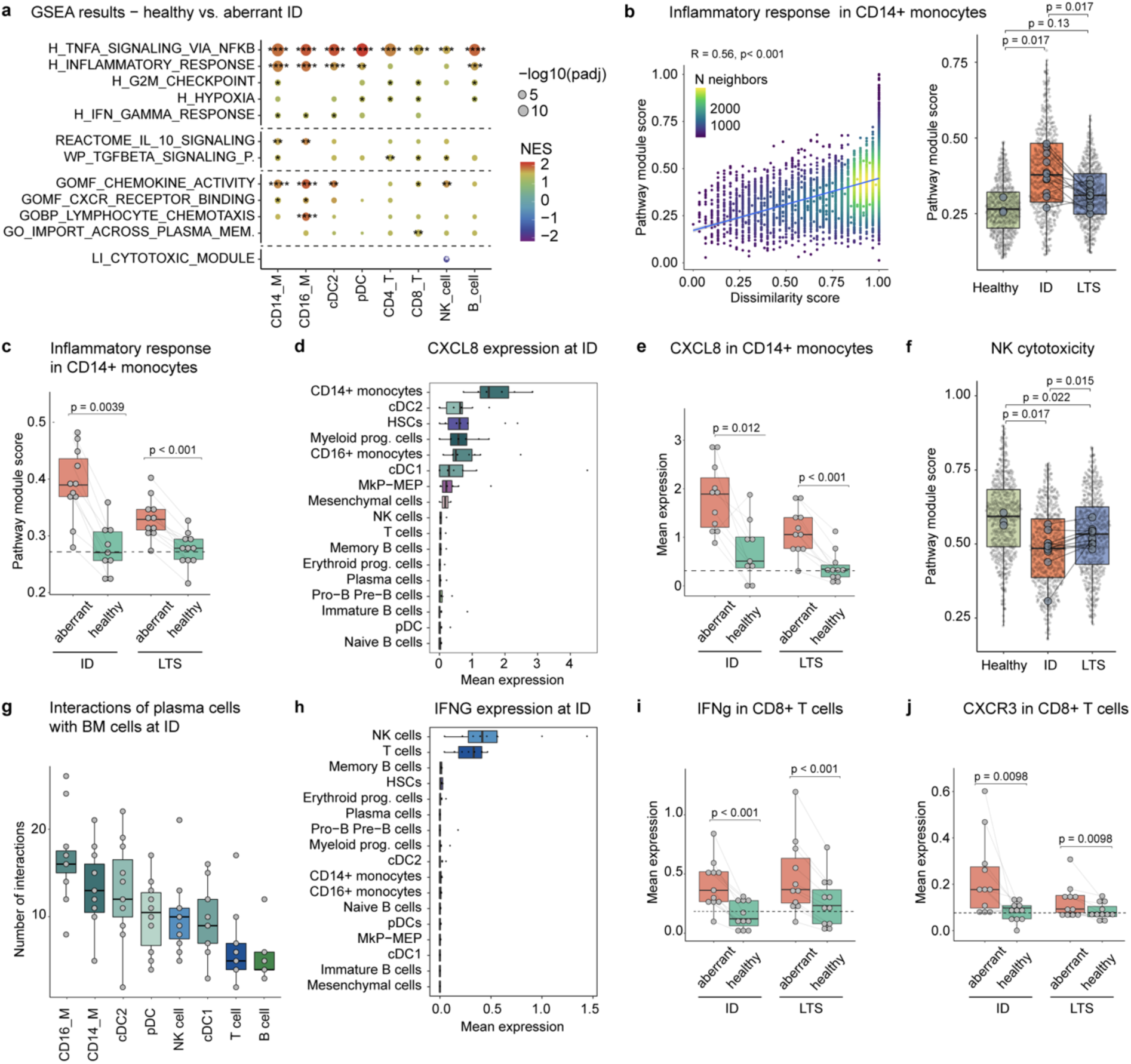
An inflammatory circuit underlies immune remodeling during active disease and long-term survival. See also extended data Figure 4. **(a)** Gene set enrichment analysis performed independently in different cell subsets from the total bone marrow and T cell datasets. Aberrant cells from patients at initial diagnosis (ID) were compared against cells from healthy donors. Selected gene sets from MSigDb Hallmark, MSigDB C2, MSigDB C5 are shown. Benjamini-Hochberg adjusted p-values are encoded by dot size, colors represent normalized enrichment scores (NES). Stars mark significant enrichment of the selected gene sets. **(b)** Left, correlation between Hallmark Inflammatory Response module score and dissimilarity score in CD14+ monocytes. The cell density is represented by color. Spearman’s Rho and the significance level of correlation are indicated. Right, distribution of the Hallmark Inflammatory Response module score by clinical group. Benjamini-Hochberg adjusted p-values from unpaired (Healthy/ID, Healthy/LTS) and paired (ID/LTS) two-sided Wilcoxon rank-sum tests are shown. **(c)** Boxplots showing Hallmark Inflammatory Response module score (see b) in CD14+ monocytes split by clinical group and cell state prediction. The dashed line highlights the mean module score within the healthy control group. Significant differences between aberrant and healthy cells were tested by comparing the respective sample means with paired two-sided Wilcoxon rank-sum tests. **(d)** Mean CXCL8 expression at initial diagnosis in the different BM cell types. **(e)** Boxplots of mean CXCL8 expression in CD14+ monocytes split by clinical group and cell state prediction. The dashed line represents the mean CXCL8 expression within CD14+ monocytes of the healthy controls. Significant differences between aberrant and healthy cells were tested by comparing the respective sample means with paired two-sided Wilcoxon rank-sum tests. **(f)** NK cytotoxicity module score in the NK cell subset summarized by clinical group. Benjamini-Hochberg adjusted p-values from unpaired (Healthy/ID, Healthy/LTS) and paired (ID/LTS) two-sided Wilcoxon rank-sum tests are shown. **(g)** Predicted number of interactions between plasma cells and immune cells from the BM at initial diagnosis derived by CellPhoneDB (see methods). **(h)** Mean interferon-gamma (IFNG) expression at initial diagnosis in different BM cell types. **(i, j)** Boxplots of IFNG (i) and CXCR3 (j) expression in CD8+ T cells split by clinical group and cell state prediction. Mean expression levels of CD8+ T cells from healthy controls are highlighted by dashed line. Significant differences between aberrant and healthy cells were tested by comparing the respective sample means with paired two-sided Wilcoxon rank-sum tests. Abbreviations: CD14/CD16_M: CD14+/CD16+ monocytes; cDC: conventional dendritic cells; pDC: plasmacytoid dendritic cells; CD4/CD8_T: CD4+/CD8+ T cells; NK: natural killer cells Box plots: center line, median; box limits, first and third quartile; whiskers, smallest/largest value no further than 1.5*IQR from corresponding hinge.

**Figure 5.**
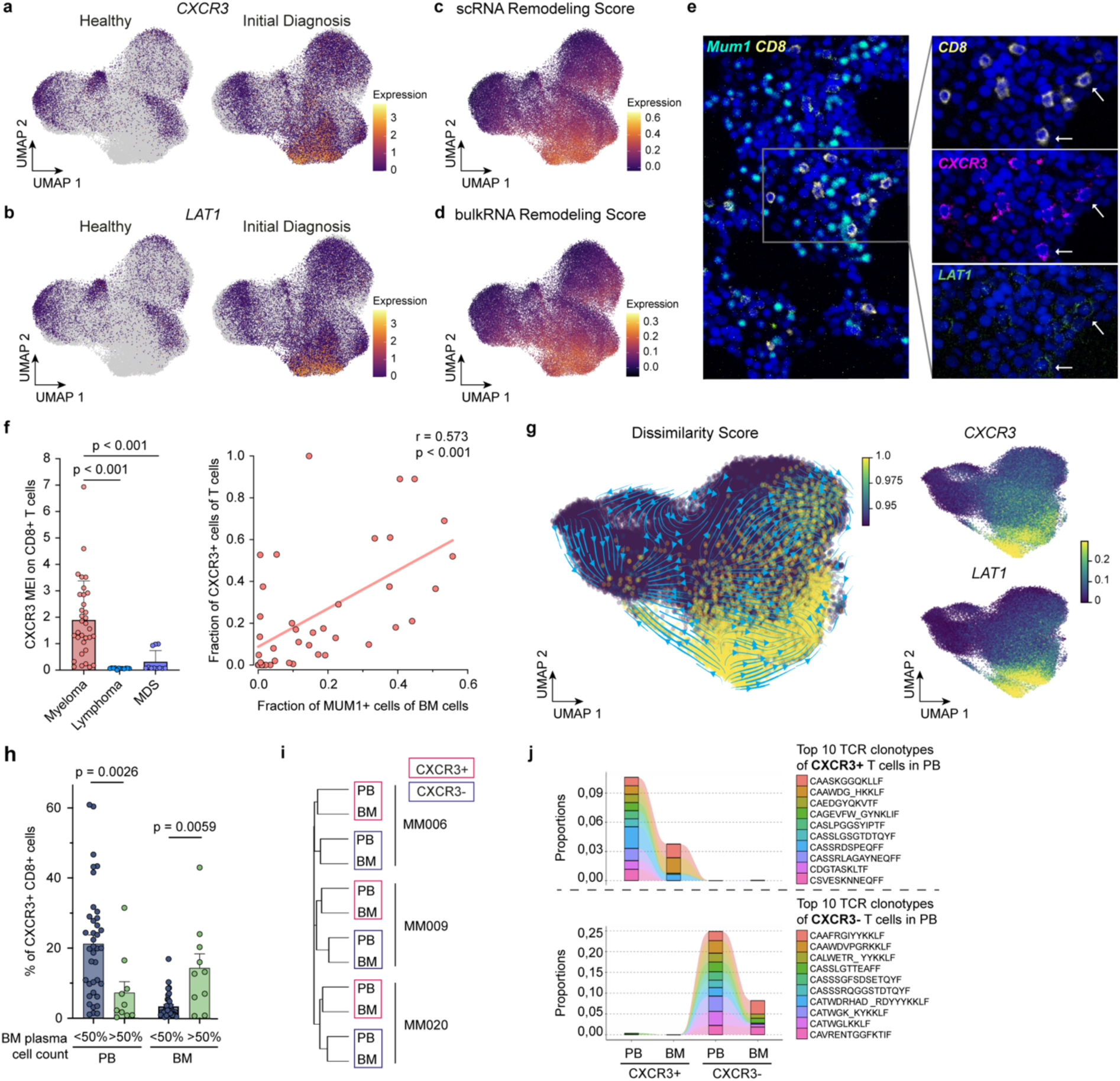
**Bone marrow infiltration of inflammatory T cells is associated with myeloma burden and serves as an accessible biomarker for disease activity.** See also extended data Figure 5. (**a**-**d**) UMAP of CD8+ T cells (derived from T cell dataset) with highlighted expression of *CXCR3* (a) and *LAT1* (b) split between cells from healthy donors and patients at initial diagnosis; and highlighted ‘scRNA Remodeling Score’ (derived from MAST analysis of scRNAseq data: healthy versus aberrant CD8+ T cells) (c) and ‘bulkRNA Remodeling Score’ (derived from DESeq2 analysis of FACS isolated CXCR3+ versus CXCR3-CD8+ T cells) (d). **(e)** Multiplex immunofluorescence images illustrating expression of *MUM1* on plasma cells as well as *CXCR3* and *LAT1* on CD8+ T cells in a representative BM area of a MM patient (examples of CD8+ T cells co-expressing *LAT1* and *CXCR3* are highlighted by arrows). **(f)** Left, *CXCR3* mean expression intensity (MEI) on BM CD8 T cells of MM patients and B Non-Hodgkin lymphoma and MDS patients as controls, Benjamini-Hochberg adjusted p-values from unpaired two-sided Wilcoxon rank-sum tests are shown; Right, Spearman correlation of the fraction of CXCR3+ T cells of all T cells with tumor burden measured by fraction of MUM1+ plasma cells in the BM. **(g)** UMAP of CD8+ T cells with highlighted velocities (arrows), dissimilarity score (yellow), and imputed *CXCR3* and *LAT1* expression. **(h)** Fraction of CXCR3+ CD8+ T cells in peripheral blood (PB) and bone marrow (BM) in patients with low to intermediate tumor burden (<50% plasma cells) versus patients with high tumor burden (>50% plasma cells) as determined by BM cytology. Significant differences between patients with low to intermediate and high tumor burden were tested by comparing the respective sample means with unpaired Wilcoxon rank sum test. **(i)** Hierarchical clustering of FACS-isolated CD8+ T cells (+/-CXCR3) from BM and PB of 3 MM patients by shared clonotypes of T cell receptor (TCR) repertoire using Jaccard index of repertoire similarity. **(j)** Clonotype tracking by representative CDR3 amino acid sequence of shared clonotypes between the top 10 most abundant TCR clonotypes from CXCR3+ (top row) and CXCR3-(bottom row) peripheral blood (PB) CD8+ T cells across CXCR3+ or CXCR3-CD8+ T cell subsets in PB and BM. Two representative patients are shown (see also extended data Figure 5). Amino acid clonotype sequences are shown as labels. Abbreviations: FACS: fluorescence activated cell sorting; BM: bone marrow; PB: peripheral blood; IF: immunofluorescence; MEI: mean expression intensity; MDS: myelodysplastic syndrome; ASCT: autologous stem cell transplantation; MFI: mean fluorescence intensity; TCR: T cell receptor.

### Bone marrow infiltration of inflammatory T cells is associated with myeloma burden and serves as an accessible biomarker for disease activity

To characterize the origin and phenotype of disease-associated remodeled immune populations, we focused on aberrant CD8+ T cells as key producers of inflammatory cytokines throughout ID and LTS. Gene expression analyses of the scRNAseq data revealed the chemokine receptor *CXCR3* and the amino acid transporter *LAT1* as accurate biomarkers for a disease-associated inflammatory CD8+ T cell state (Fig.5a-b). To further assess the value of surface *CXCR3* expression as a marker for myeloma-associated CD8+ T cells, we subjected BM *CXCR3+* and *CXCR3-* CD8+ T cells from an independent cohort of 7 MM patients to bulk RNA-sequencing (Extended data Fig. 5b). Importantly, scRNAseq-derived *CXCR3* expression was highly overlapping with both, the single-cell derived gene signature defining aberrant CD8+ T cells (Fig. 5c) and the bulk RNAseq-derived gene signature for CXCR3+ T cells within the BM (Fig. 5d). This confirms the specificity of surface *CXCR3* as biomarker for remodeled inflammatory T cells.

Next, we performed multiplex immunofluorescence stainings on BM biopsies and confirmed the co-expression of *CXCR3* and *LAT1* on CD8+ T cells in MM patients (Fig. 5e). Importantly, the mean expression intensities of *CXCR3* as well as *LAT1* in CD8+ T cells were highly elevated in MM patients compared to B cell Non-Hodgkin lymphoma and MDS control cohorts, confirming the specific enrichment of aberrant inflammatory CD8+ T cells in MM (Fig. 5f, Extended data Fig. 5c). Notably, the fraction of detected aberrant inflammatory CD8+ T cells positively correlated with the number of *MUM1*+ plasma cells, suggesting that *LAT1* and *CXCR3* can serve as a biomarker for both tumor load and associated remodeling of the BM immune microenvironment (Fig. 5f, Extended data Fig. 5c).

To explore the origin of remodeled CD8+ T cells, we determined RNA velocities to predict the future cell state based on ratios of spliced to unspliced mRNAs (see methods). As reported in previous studies, this analysis revealed the transient and connected states of the main T cells subsets [26] (Fig. 5g). However, the cluster comprising aberrant inflammatory T cells marked by *LAT1* and *CXCR3* expression and a high dissimilarity score appeared disconnected to the cluster harboring the main homeostatic BM-resident T cell subsets (Fig. 5g). As described above, *CXCR3* is a chemokine receptor mediating migration towards the chemoattractants *CXCL9/10/11,* which are synthesized at increased levels in the BM upon MM (Extended data Fig. 4k,l). These observations point towards a chemokine-mediated infiltration of inflammatory T cells from the periphery to the BM. To further explore this, we quantified the *CXCR3* expression on CD8+ T cells of paired BM and peripheral blood (PB) samples from 48 MM patients via flow cytometry (Extended data Fig. 5b). In line with our hypothesis, in patients with low tumor burden (<50%), *CXCR3*+ T cells were mainly present in the peripheral blood and not in the BM (Fig. 5h). In contrast, in patients with high tumor burden (>50%) the number of *CXCR3*+ T cells decreased in PB, while an increased number of *CXCR3+* T cells was observed in the BM, suggesting a tumor-load dependent migration of inflammatory T cells to the BM.

To further validate this finding, we isolated *CXCR3*+ and *CXCR3*-CD8+ T cells from paired PB and BM samples of newly diagnosed MM patients and performed bulk RNA-sequencing, followed by mapping the T cell receptor (TCR) repertoire (Extended data Fig. 5b). While we did not observe any indication for clonal expansion of inflammatory *CXCR3*+ CD8+ T cells (Extended data Fig. 5d), hierarchical clustering based on TCR repertoire information revealed a striking overlap of the *CXCR3*+ fractions from PB and BM for each patient, indicating a close relation between remodeled CD8+ T cells in the BM with *CXCR3*+ CD8+ T cells in PB (Fig.5i, Extended data Fig. 5e). In line with this, the clonotypes of the top 10 clones in *CXCR3*+ T cells from the PB showed a high overlap with the top clonotypes in *CXCR3*+ T cells from BM fraction but not with their *CXCR3*-negative counterparts, demonstrating a disease-associated infiltration of inflammatory T cells from the periphery to the BM (Fig. 5j, Extended data Fig. 5f).

Together, our data reveal that upon MM disease activity, inflammatory CD8+ T cells are recruited to BM where they serve as key players in the establishment and maintenance of the sustained inflammatory BM remodeling at ID and LTS (Extended data Fig. 5a). BM infiltration by inflammatory T cells is associated with myeloma burden and serves as an accessible biomarker for disease activity that can be measured both in the BM and the peripheral blood.

### Immune remodeling in LTS patients is associated with future disease resurgence and impaired immune function even in the absence of measurable disease

We next investigated the underlying causes of sustained immune alterations in MM LTS patients. A sustained persistence of malignant plasma cells or a resurgence of the disease may trigger immune perturbations. To investigate this hypothesis, we compared the degree of immune cell remodeling as measured by DA-seq-based prediction scores to the fraction of malignant plasma cells present within the overall plasma cell pool for each patient. Indeed, microenvironmental immune remodeling was associated with the proportion of malignant plasma cells present in the bone marrow (Fig. 6a). This finding indicated disease burden as an important factor for sustained immune perturbation in the BM. In line with this, the degree of immune remodeling and the activity of pathways that are upregulated during the full-blown disease gradually increased from healthy donors to LTS patients with CR to patients who experienced relapse from CR (termed non-CR patients), confirming a tumor-burden associated remodeling (Fig. 6b,c). This step-wise remodeling was consistently observed in the CD8+ T cell compartment as well as in other immune compartments (Extended data Fig. 6a-f). Accordingly, *CXCR3* expression on CD8+ T cells, which we have identified as a surrogate marker for disease activity, was correlated with the fraction of malignant plasma cells present in the BM (Fig. 6d).

**Figure 6.**
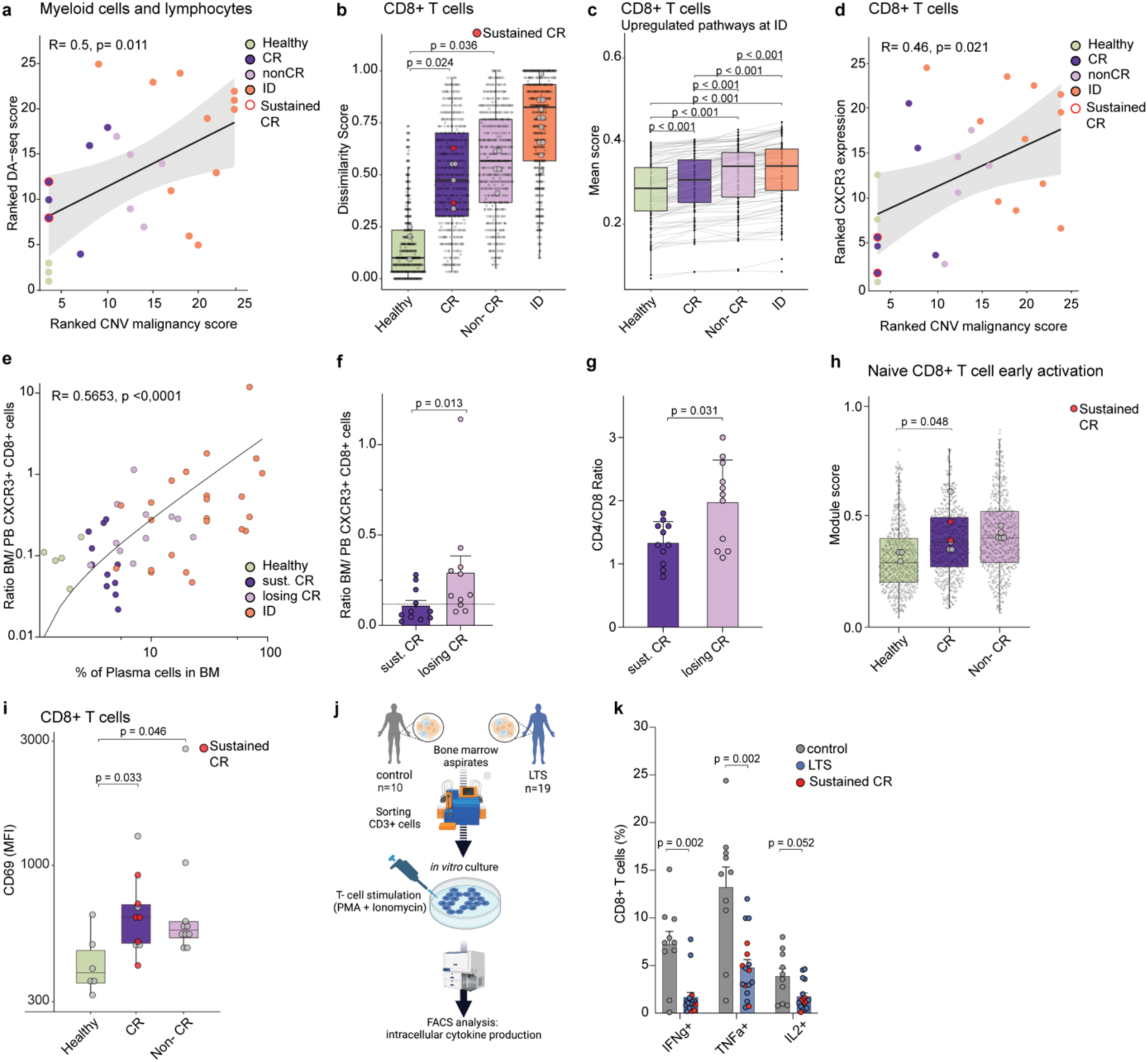
Immune remodeling in LTS patients is associated with future disease resurgence and defective immune function even in the absence of measurable disease. Also see extended data. **Figure 6**. **(a)** Correlation of the malignant plasma cell fraction of all plasma cells (CNV-based malignancy score) and the degree of remodeling (as quantified by the mean DA-seq score). Scores are ranked by the mean expression. Each dot represents the ranked DA-seq score across immune cells and the corresponding CNV malignancy score. Spearman’s Rho and the significance level of correlation are indicated. Colors represent clinical groups. **(b)** Distribution of the dissimilarity score (small dots) by clinical group with LTS patients split into patients with complete remission (CR) and patients with biochemical progression (non-CR) patients within the CD8+ T compartment. Large dots indicate sample means. Sustained CR patients are highlighted in red. Benjamini-Hochberg adjusted p-values from unpaired two-sided Wilcoxon rank-sum tests are shown. **(c)** Mean module scores of top 100 upregulated pathways (ID vs. Healthy) between clinical groups in CD8+ T cells. Benjamini-Hochberg adjusted p-values from paired one-tailed Wilcoxon signed rank test are highlighted. **(d)** Correlation between mean CXCR3 expression in CD8+ T cells per patient and the malignant plasma cell fraction of all plasma cells per patient (CNV-based malignancy score); Spearman’s Rho and the significance level of correlation are indicated. **(e)** Correlation between the ratio of BM to PB CXCR3+ CD8+ T cells as measured by flow cytometry and cytological plasma cell count in the BM per sample. Individual patients are highlighted as dots. Clinical groups are highlighted by color. Spearman’s Rho and the significance level of correlation are indicated. **(f)** Comparison of the BM to PB ratio of CXCR3+ CD8+ T cells between patients that experienced sustained CR or lost the CR state during a 4-year clinical follow up (losing CR) quantified by flow cytometry. Individual patients are highlighted as dots. Mean ratio of healthy controls is highlighted by dashed line. Significance was tested by unpaired Wilcoxon rank sum test. **(g)** Comparison of CD4+ to CD8+ T cell ratios between patients with sustained CR versus patients losing CR within peripheral blood quantified by flow cytometry. Individual patients are highlighted as dots. Significance is shown for unpaired Wilcoxon rank sum test. **(h)** Boxplot of module scores for the CD8 early activation gene signature from [27] in naive CD8+ T cells. Data is summarized and statistically tested as described in b. **(i)** Mean CD69 expression in CD8+ T cells measured by flow cytometry and compared between healthy controls, CR and Non-CR patients that experienced LTS. Patients in sustained CR are highlighted in red. Significance was tested using two-sided, unpaired Wilcoxon rank sum test and corrected according to Benjamini-Hochberg. **(j)** Study design scheme of the *in vitro* T cell cytokine assay. CD3+ T cells from 19 LTS patients and 10 controls were isolated by FACS and stimulated with PMA and Ionomycin. Intracellular cytokine production was assessed by flow cytometry. **(k)** Quantification of intracellular CD3+ T cell cytokine expression of IFNg, TNFa and IL2 from LTS patients and controls as determined by flow cytometry. Patients in sustained CR are highlighted in red. Significant differences between controls and LTS patients were tested by comparing the respective sample means with unpaired Wilcoxon rank sum test. Abbreviations: ID: initial diagnosis; LTS: long-term survival; CR: complete remission; BM: bone marrow; PB: peripheral blood; FACS: fluorescence activated cell sorting.

Clinical follow-up of LTS patients over the next four years after sample acquisition revealed that even patients that had been in CR for over a decade may exhibit signs of relapse while others experience a sustained CR (Extended data Fig. 6g). Importantly, at time of sample acquisition, the ratio of BM to peripheral blood *CXCR3*+ CD8+ T cells determined by flow cytometry in a larger validation cohort (5 healthy donors, 24 LTS patients and 23 MM patients at ID) gradually increased from healthy donors via sustained CR and patients losing CR to ID patients, reflecting the respective disease burden of the different clinical states (Fig. 6e). Moreover, CR patients that will lose their CR within the next four years, displayed a significantly higher BM to blood *CXCR3*+ CD8+ T ratio if compared to patients that will remain in sustained CR (Fig. 6f). In line with an increased CD8+ T cell infiltration into the BM, an increased CD4+ to CD8+ T cell ratio in the blood was also associated with future relapse from CR during LTS (Fig. 6g). These data demonstrate that blood measurements can be used as accessible biomarkers to track environmental perturbations in the BM associated with future relapse.

Collectively, our findings suggest that perturbations observed in the immune microenvironment are in part triggered by the sustained presence of malignant plasma cells or resurgence of the disease in LTS patients. However, even in the absence of any measurable disease activity at the time of sample acquisition at LTS, as well as during a 4-year follow-up (sustained CR), the immune remodeling was still apparent, suggesting long-lasting, irreversible disease or therapy-associated immunological changes (Fig. 6a-c). In particular, the naïve CD8+ T cell compartment of LTS patients showed higher expression of an ‘early T cell activation signature’ even in the absence of any measurable disease activity, pointing towards a chronic pre-activatory state (Fig. 6h) [27]. A gene within this ‘early activation signature’ was the well-studied surface marker CD69, expressed on activated human T cells [28]. In line with our previous findings, CD8+ T cells from patients in CR and ‘sustained’ CR displayed increased CD69 surface protein levels confirming the persistent long-term imprint on CD8+ T cells in the absence of disease activity (Fig. 6i). To assess whether these sustained aberrations also translate into changed T cell functionality, we measured the capacity of T cells from LTS patients to produce cytokines upon T cell activation. For this, MACS-sorted CD3+ T cells were stimulated with PMA and Ionomycin, and intracellular cytokine production (TNFa, IFNg, IL2) was measured as a surrogate parameter for T cell functionality (Fig. 6j). Notably, stimulated T cells from patients in LTS produced significant lower amounts of all measured cytokines compared to control samples from healthy controls and early stage MM patients (Fig. 6k). Of note, the impaired T cell functionality was also observed in patients with no measurable disease activity and sustained CR, suggesting a sustained immunological scarring in LTS patients. Together, our study reveals persistent immune remodeling and impaired T cell functionality upon LTS even in patients with sustained CR in the absence of any measurable disease activity.

## Discussion

The long-term consequences of cancer and cancer therapy on the immune system remain poorly understood. In this study, we have comprehensively investigated the immune ecosystem in MM LTS patients years to decades after successful first line therapy. We uncovered that MM long-term survivors display sustained immune alterations that are associated with the resurgence of the disease and correlated with disease activity. These disease-associated immune alterations are mediated by an inflammatory circuit driven by a tumor load-dependent infiltration of inflammatory T cells into the bone marrow. However, even in the absence of any measurable disease activity for years to decades, long-term alterations in the bone marrow ecosystem associated with defective immunity were observed.

Previous studies on immune reconstitution after exposure to cancer or cancer therapy, including autologous stem cell transplantation, focused on the short-term impact. For example, Boekhorst et al [29] and Schlenke et al [30] investigated the reconstitution of the T cell compartment in a mixed cohort of different hematological, as well as solid tumor patients. Both studies did not observe any signs of functional impairment in T cells from PB as measured by standard flow cytometry phenotyping. However, MM patients were underrepresented in both study cohorts. In a study focusing on short-term consequences of auto-SCT in MM, an impaired cytokine production of the T cell compartment was observed, concluding that the complete recovery of the immune system might require more time [31]. However, our study reveals long-term sustained molecular changes in the immune microenvironment, even in MM patients that were considered functionally cured, suggesting an irreversible immunological scarring, as previously described in infectious diseases [6, 7]. While our study focused on transcriptomic and immunological changes in LTS patients, a recent study identified clonal hematopoiesis as a common event upon long-term survival of pediatric cancers [32]. In a subset of Hodgkin Lymphoma survivors, therapy-related STAT3 mutations were detected that potentially also impact on T cell biology. While our data support a non-genomic mechanism of sustained changes of the immune system in MM LTS, we cannot exclude that also genomic aberrations may contribute to some of the irreversible phenotypes we observed.

Our study revealed a tumor load dependent inflammatory circuit in MM with the release of CXCL9/10/11 from myeloid cells causing the migration of CXCR3+ inflammatory T cells from the periphery to the BM, in line with previous reports in the context of cancer and vaccinations [33-35]. Inflammatory T cells and NK cells in turn act as major drivers for IFNg-mediated BM changes in a self-propelling circuit. This inflammatory circuit is initiated at ID and maintained in a subset of immune cells during LTS. Importantly, we demonstrate that disease-associated T cell trafficking can be used to track and reliably predict the re-initiation of the disease in LTS patients in the bone marrow by analyzing CXCR3 expression on CD8+ T cells and the immune composition (CD4/CD8 T cell ratio) in the peripheral blood. This highlights how disease associated changes in the microenvironment might be used in combination with MRD detection methods to predict resurgence of disease activity. While the detailed contributions of T cell migration to anti-cancer immunity remains to be investigated, targeting the introduced inflammatory circuit may offer potential avenues for new therapeutic strategies [36, 37]. Of note, our study included paired samples of patients experiencing long-term remission after a single therapy line in the absence of any maintenance therapy for years. Due to continuous maintenance therapy as the new standard of care, this patient cohort is not recruitable nowadays and thus displays a unique selection of patients to study the long-term consequences of cancer and cancer therapy in absence of potential biases associated with additional therapies.

Together, our study provides detailed insights into the molecular and cellular bone marrow ecosystem of MM long-term survivors, thereby revealing reversible and irreversible disease-and therapy-associated alterations of the immune compartment which can serve as diagnostic and predictive tools.

## Acknowledgments

We would like to thank members of the Haas, Hundemer and Brors laboratories for helpful discussions. Moreover, we thank members of the DKFZ Single cell open lab and the DKFZ flow cytometry for support. We would like to thank A. Mahmoud for an initial analysis of the single-cell RNA-seq dataset as part of his PhD thesis. This study was supported by a grant from the Black Swan Research Initiative of the International Myeloma Foundation, the German Bundesministerium für Bildung und Forschung (BMBF) through the Juniorverbund in der Systemmedizin ‘LeukoSyStem’ (FKZ 01ZX1911D) and the Baden-Württemberg Stiftung (FKZ MET-ID43). Contribution by R.L was supported by the DKFZ Clinician Scientist Program funded by the Dieter Morszeck Foundation.

## Author Contributions

R.L., S.H., M.H. and H.G. conceived the study. R.L. performed the single-cell RNA sequencing experiments with help from T.B. and S.H.. R.L., M.B. and M.A. performed the experimental validations and functional experiments with help from M.H., D.V. and C.W.. F.G. and M.S. conducted the majority of bioinformatics analyses with conceptional input from R.L., S.H., M.H., B.B. and C.I.. L.J.-S., M.B., S.Y., N.B., A.S., C.I. and G.S. performed additional bioinformatics analyses. S.H. and M.H. supervised the experimental work. C.I. and B.B. supervised the bioinformatics analyses with conceptional input from S.H. and D.H.. D.V., T.B., D.H., B.G., N.W., M.S.R, C.W, A.T., H.G. and C M.-T. provided clinical samples and conceptional input on data interpretation. R.L., S.H., F.G., M.S., M.H., N.B., L.J-S., S.Y, M.B., A.S. and C.I. wrote the manuscript and prepared figures. All authors have carefully read the manuscript.

## Materials and Methods

### Human samples

#### Ethics approval and consent to participate

BM samples from healthy and diseased donors were obtained at Heidelberg University hospital after informed written consent using ethic application numbers S-480/2011 and S-052/2022. BM aspirates were collected from iliac crest. Healthy BM donors received financial compensation in some cases. For BM, mononuclear cells (BMMCs) were isolated by Ficoll (GE Healthcare) density gradient centrifugation and stored in liquid nitrogen until further use. All experiments involving human samples were approved by the ethics committee of the Heidelberg University hospital and were in accordance with the Declaration of Helsinki.

### Flow cytometry

#### MRD analysis

Flow cytometry for detection of minimal residual disease (MRD) in fresh human BM samples was performed according to the highly standardized flow cytometry approach developed and described by the Spanish Myeloma Collaborative Group using a commercially available EuroFlow 8-color 2-tube MM MRD Kit (Cytognos, Salamanca, Spain) [38]. Tube one contained multiepitope CD38-FITC, CD56-PE (clone C5.9, CD45-PerCP-Cyanine5.5 (clone EO1), CD19-PE-Cyanine7 (clone 19-1), CD117-APC (clone 104D2) and CD81-APC-C750 (clone M38) antibodies. Tube two contained multiepitope CD38-FITC, CD56-PE (clone C5.9), CD45-PerCP-Cyanine5.5 (clone EO1), CD19-PE-Cyanine7 (clone 19-1), cytoplasmic polyclonal immunoglobulin (Ig) κ-APC goat and cytoplasmic polyclonal Igλ-APC-C750 antibodies. Drop-in CD27 Brilliant Violet 510 (clone O323, Biolegend, San Diego, USA) and CD138 Brilliant Violet 421 (clone MI15, BD, Heidelberg, Germany) antibodies were added to tubes one and two. Measurements were performed on a cell analyzer (BD, Heidelberg, Germany) after implementation of the EuroFlow Standard Operating Protocol for Instrument Setup and Compensation in FACSDiva (BD Biosciences, San Jose, CA, USA). Final data analysis was performed in Infinicyt 2.0 (Cytognos, Salamanca, Spain). An automated gating and identification tool (Cytognos, Salamanca, Spain) was used to support the identification of MM cells. Plasma cells were identified based on the co-expression of CD38 and CD138 antigens. An aberrant plasma cell expression profile was defined as CD45-low/negative, CD56-positive, CD19-negative and light chain-restricted.

#### Flow cytometry of cryopreserved BM samples

Human BM samples were thawed in a water bath at 37 °C and transferred dropwise into RPMI-1640 10% FCS. Cells were centrifuged for 5 min at 350 rpm and washed once with RPMI-1640 10% FCS. Cells were resuspended in FACS buffer (FB) (PBS 5% FCS 0.5 mM EDTA) containing different antibody cocktails (see below) and FcR blocking reagent (Miltenyi) and incubated for 15 min at 4 °C.

For analysis of CXCR3 expression on CD8+ T cells across different clinical groups, cells were stained with CD8, CD3, CD45, CD4, CD194, CD196, CD152, CCR10 surface antibodies. For analysis of CD69 expression on CD8+ T cells, cells were stained with CD8, CD97, CD4, CXCR4, CD26, CD45RO, CD6, CD69, CD98, CD29, CXCR3, CCR7 and CD3 surface antibodies.

After washing with FB, all experiments were measured on BD FACSFortessa flow cytometer, equipped with 5 lasers, or BD FACSLyric flow Cytometer, equipped with three lasers.

### Single-cell RNA sequencing data

#### BM preparation, staining and sorting for gene expression analysis

Human BM samples were thawed in a water bath at 37 °C and transferred dropwise into RPMI-1640 10% FCS. Cells were centrifuged for 5 min at 350 rpm and washed once with RPMI-1640 10% FCS, followed by resuspension in FACS buffer (FB) (PBS 5% FCS 0.5 mM EDTA) containing CD45-PE and CD3-APC and FcR blocking reagent (Miltenyi) and incubation for 15 min at 4 °C. Cells were washed with FB. To exclude debris and ensure that actual cells were sorted for droplet-based scRNAseq, cells were stained with a DNA dye (Vybrant DyeCycle Violet, Thermo Fisher Scientific). For this purpose, 2.5 µl ml−1 Vybrant dye in cell suspension medium was incubated with 3 × 10^6 cells at 37 °C for 20 min in a water bath. Following the incubation, the cells were placed on ice and were sorted immediately for each experiment into 15 µl PBS containing 2% fetal bovine serum. For sorting of total BM cells, single, live cells were gated and sorted. For sorting of T cells CD45+ CD3+ cells were gated and sorted. Cells were sorted using a FACSAria Fusion or FACSAria II equipped with 100 µm nozzles respectively. Sorted cell numbers were confirmed using a LUNA automated cell counter (Logos Biosystems). A volume of 33.8 µl of the cell suspension was used as input without further dilution or processing, with final concentrations around 300 cells per µl.

#### Single-cell RNA sequencing and data preprocessing

Single-cell RNA sequencing libraries of BMMCs form healthy controls and MM patients were generated using 10x Genomics single-cell RNAseq technology (Chromium Single Cell 3’ Solution v2) according to the manufacturer’s protocol and sequenced on an Illumina HiSeq4000 (paired end, 26 and 74 bp). Upon sequencing, FASTQ files were processed and aligned to the human reference genome GRCh38 (GENCODE v32) using the standard Cellranger pipeline (10x Genomics, v4.0).

### scRNA-seq data analysis

All analyses were performed in R (v4.0.0). The output from the Cellranger pipeline was combined into one count matrix and further processed and analyzed using the Seurat framework (v4.0.1, [39]). Parameters are indicated when non-default settings for a specific function were used.

#### Quality control of BM scRNA-seq data

Cells were excluded for downstream analysis if they where of low quality (< 200 UMIs, < 400 detected features, > 10% mitochondrial counts), were identified as doublets by library size and expressed features (> 40.000 UMIs, > 6.000 detected features), or if they did not express cell-type-or -state-specific genes. In addition, decontX()from the R package celda (v1.4.7, [40]) was used to estimate and remove contaminating ambient RNA.

#### Dimensionality reduction and clustering of BM scRNAseq-data

Gene counts were log-normalized and the top 2000 variable features were identified and scaled using default parameters of FindVariableFeatures() and ScaleData(). Dimensionality reduction of the scaled data was performed by principal component analysis (PCA). The top 50 PCs were then used to build a shared nearest neighbor graph (SNN, FindNeighbors(dims=1:50)) for Louvain clustering (FindClusters(resolution=0.7)) and uniform manifold approximation and projection (RunUMAP(Dims=1:50)) of the data in two-dimensional space. Final cluster resolution and annotation was defined by evaluating known marker genes. Clusters with overlapping gene signatures were merged to reach overall cell-type resolution (*MetaClusters*).

In order to achieve a more fine-granular filtering and annotation, each cell-type (*MetaCluster*) was subsetted and count matrices were separately processed again from variable feature selection and re-scaling to dimensionality reduction by PCA and subsequent clustering and UMAP representation. Clusters with contaminating gene expression profiles, or aberrantly high mitochondrial and low housekeeping gene expression were considered as doublets, or low quality, respectively and removed. Final cell annotation was then transferred back to the global BM count matrix. In addition, cells from patients treated with maintenance and induction therapy were removed.

#### Copy number analysis

Single-cell copy number analysis was performed using infercnv (v1.6.0, [41]) with JAGS (v4.3.0, [41]) with JAGS (v4.3.0, [42]). First, we generated a gene ordering file using a Python script provided by the infercnv developers (https://github.com/broadinstitute/infercnv/blob/master/scripts/gtf_to_position_file.py, 21 Apr 2021) and excluded all genes that were not part of this file. We only considered chromosomes 1-22 and, in order to avoid artefacts due to differential immunoglobulin gene expression, excluded all genes starting with “*IGH*”, “*IGL*” or “*IGK*”. The actual inferCNV analysis was performed separately for the plasma cells from each patient and utilized non-normalized decontX-corrected expression values. Plasma cell from the three healthy donors were used as reference cells. We disabled the filtering threshold regarding counts per cell and used the arguments “cutoff = 0.1”, “cluster_by_groups = TRUE”, “cluster_references = FALSE”, “analysis_mode = ‘subclusters’. “tumor_subcluster_pval = 0.05”. “denoise = TRUE”, “noise_logistic = TRUE”, “HMM = TRUE”, “HMM_type = ‘i6’” and “num_threads = 1” within infercnv’s function run(). Subsequently, we manually annotated the detected sub-populations as “healthy”, “malignant” or “unclear” based on the denoised infercnv results. We additionally determined the major immunoglobulin light chain expressed by malignant cells in a patient-wise fashion by inspecting the expression of the corresponding genes (*IGKC, IGLC1-7*). Afterwards, we refined the malignancy annotation to reduce the number of cells that were wrongly classified as malignant. To this end, we compared immunoglobulin light chain gene expression (decontX-corrected and normalised) in each putatively malignant cell with the corresponding mean expression in its sub-population. If the expression of the patient-specific major light chain gene was less than half of the corresponding mean expression in the corresponding sub-population and the expression of another light chain gene was above 1.5 times the corresponding mean expression in the corresponding sub-population, a cell’s classification was forced to “healthy”. Copy number heat maps were generated using ComplexHeatmap (v2.6.2, [43]), circlize (v0.4.13, [44]), scales (v1.1.1, [45]), magick (v2.7.3, [46]) and imagemagick (v6.9.12, [47]). Only cells from samples that were not obtained during induction and maintenance treatment are displayed.

#### scRNA-seq quality control of T cell data

Cells were kept in the dataset if they had between 500 - 20000 UMIs, between 300 - 4000 detected features and less than 10% mitochondrial reads. Clusters of contaminating cells including myeloid cells, erythroid progenitors and plasmablasts were identified based on expression of cell type-specific marker genes. Subsequently, `decontX()` from the R package celda (v1.4.7, [40]) was applied on the count matrix to account for cross-contaminating reads using the contaminating cell types and remaining T cells as cluster labels. The final Seurat object was filtered to maintain only T cells and the decontX matrix was used for all subsequent analyses.

#### Classification of T cell subsets

A reference dataset was generated from the T cell dataset by annotating cells based on the normalised decontX matrix (NormalizeData): CD4: CD4 > 1.5 & CD8A == 0 & CD8B == 0 & TRDC ==0 CD8: (CD8A > 1.5 | CD8B > 1.5) & CD4 == 0 & TRDC ==0 gdT: TRDC > 1.5 & CD8A == 0 & CD8B == 0 & CD4==0 For each of these T cell subsets, dimensionality reduction was performed ((`NormalizeData()`, `FindVariableFeatures(nfeatures=1000)`, `ScaleData()`, `RunPCA()`) and cells were clustered to define the main cell states (`FindNeighbours(reduction=’pca’.dims=1:20).`

`FindClusters(resolution=0.4)`). The subsets were then merged back into a combined reference dataset to annotate the complete T cell dataset with SingleR (v1.2.4, [48]) taking “pruned.labels” output to split the T cell Seurat object into CD4, CD8 or gdT cell subsets for further analyses.

#### CD8 subset analysis

Dimensionality reduction and clustering was re-run (as above, except `RunUMAP(dims=1:20)`, `FindClusters(resolution=0.5)`) as final filtering step excluding a cluster specific for cycling cells and then repeated to obtain a final version (as before, except `FindClusters(resolution=0.45)`). Clusters were annotated to CD8+ T cell states based on the module score expression for custom gene signatures, which was added for each cell with `AddModuleScore()`: naive (genes: CCR7, TCF7, LEF1, SELL; cluster: 1), effector/central memory (genes: GPR183, CCR7, SELL, IL7R, CD27, CD28, GZMA, CCL5, S1PR1, GZMK, CXCR4, CXCR3, CD44; clusters: 2, 3, 5, 7), cytotoxic (genes: EOMES, TBX21, GZMB, PRF1, FASLG, GZMH, GZMA; cluster: 4). Additionally, cluster 6 was annotated as KLRB1+ T cells based on the high expression level of the corresponding gene.

#### CD4 subset analysis

Similar to the CD8+ T cell dataset, cells were projected into a low dimensional space and grouped using graph-based clustering (as before, except `FindVariableFeatures(nfeatures=3000)`, `FindClusters(resolution=0.45)`).

#### Differential Abundance Analysis

Changes in the composition of the BM microenvironment between the clinical states were evaluated by log2fold-change difference of each patient’s cell type fraction from the corresponding healthy control’s mean fraction. For differential compositional analysis (DPA) of the immune compartment, plasma cells and erythroid progenitors were excluded prior to calculating each patient’s composition per clinical group, which were tested for significance using unpaired Wilcoxon rank sum test.

For cluster-independent differential abundance analysis, DA-seq was performed [49]. The tool computes a multiscale score for each cell based on the k-nearest-neighbourhood for k between 50 and 500. Cells with a multiscale score > 0.95 and < - 0.95 were considered as differential abundant. Subsequently, a logistic regression classifier was trained on the multiscale score to obtain the differential abundant clusters which were visualized on the UMAP. A continuous DA-seq score was calculated by subtracting scaled module scores (AddModuleScore()) for significantly up-and downregulated genes in differentially abundant cells.

#### Dissimilarity analysis and aberrant cell classification

To determine and quantify whether a cell is transcriptionally more similar to healthy cells or to perturbed counterparts in the disease state, we introduce a ‘dissimilarity score’. It requires condition labels *i* (in our case “Healthy” and “ID”), sample labels *j* and a data matrix *X*. The analysis was performed per cell type to account for cell type-specific transcriptional differences. By default, we chose PCA coordinates of *n* dimensions as dimension-reduced representation of our data, where *n* was assessed by prior MetaCluster analysis. Cells were divided by condition and further sampled to adjust for equal group sizes. We computed the k = 30 nearest neighbors using the *FNN* package (v.1.1.3) to look at the condition distribution for each cell in the dataset. Dissimilarity was quantified by summing up the neighbors per condition with higher values meaning more neighboring cells from the diseased state (ID) as compared to healthy. To adjust for sampling effects, this process was iterated 100 times with changing seeds. Each cell is assigned the median dissimilarity and the final score is scaled between 0 and 1 between all conditions.

To allow group-wise comparisons between ‘healthy-like’ cells and most dissimilar, i.e. ‘aberrant-like’ cells among the clinical states, we used the automatic machine learning software H2O autoML [50]. Initially, each cell was given a ‘state’ label (‘healthy-like’ or ‘aberrant-like’) based on the combination of the ‘clinical state’ (‘Healthy’ or ‘ID’) and the ‘dissimilarity score’. The underlying ‘dissimilarity score’ threshold was defined as 99% of all cells from the healthy controls being labeled ‘healthy-like’, and applying this threshold on all patients’ cells. Then, top 500 to 1000 variable genes were computed for each cell population (see table 2) using *Seurat’s* `FindVariableFeatures()`. To train and validate the models, training (80%) and test (20%) datasets were generated for each cell population using the `createDataPartition()` function from the *caret* package (v.6.0-91, [51]). To have sufficient numbers of healthy plasma cells for model training and validation, healthy plasma cells from the ‘Human Cell Atlas’ [52] were integrated with our dataset applying the Scanorama algorithm with default parameters on all features [53]. The partitioned datasets were then converted to H2O objects using the H2O library (H2O R version: 3.36.0.3. H2O cluster version: 3.36.0.3). The function `h2o.automl()` was used for the model training process using the train dataset and top n variable genes (500 or 1000) as input. Following parameters were set: max_models = 80 (which computes 82 models due to including the two Stacked Ensembles as default), max_runtime_secs_per_model = 7200, stopping_rounds = 5 and nfolds= 50 or nfolds = 5 (depending on dataset size). Moreover, a seed was set to ensure result reproducibility.

The top leader model (see table 2) was selected and used for label prediction on the respective test dataset. To assess label prediction accuracy for each model, a confusion matrix was generated and the F1 score calculated using *caret’s* `confusionMatrix()` function. The respective leader model was then used for classification and label prediction. After running `h2o.predict()`, additional filtering thresholds were applied (p0 >= 0.66 and p1 >= 0.66) on the internal probability values to differentiate between clearly defined (p0 >= 0.66 and p1 >= 0.66) and non-defined cells.

#### Differential gene expression analysis

Differential gene expression analyses were computed using a two-part generalized linear model implemented in MAST (v1.18.0, [54]). The Hurdle model in MAST considers the bimodal expression distributions of single-cell data having either a strong gene expression or zero values (zero-inflation). Normalized decontX corrected data of the whole human bone marrow or without the cells of ID were used as input. Genes with less than 10% expression across all libraries were filtered out. For the remaining genes the hurdle model using the patients, the cell state and CR status was fitted using the MAST function `zlm()`. The obtained coefficients for each variance-covariance and gene were reported with `summary()`.

#### Gene set enrichment analysis

Gmt files containing gene set collections were obtained from Molecular Signatures Database (c2.cp.v7.4.symbols.gmt, c5.all.v7.4.symbols.gmt, h.all.v7.4.symbols.gmt, [55],[56]). To search for enriched terms of cells from patients at initial diagnosis being classified as ‘aberrant’ compared to ‘healthy’ cells from healthy donors, their average log2 fold-change among all genes was calculated. Subsequently, genes were sorted by their average log2 fold-change and used for multilevel GSEA with the fgsea R package (v1.14.0, [57]). Results were filtered for padj < 0.05 and sorted by their normalized enrichment score (NES). Significantly enriched gene sets of interest were further evaluated by calculating a module score for the corresponding gene signature, or for specified leading-edge genes in each cell using AddModuleScore()in Seurat and comparing these modules in cell types of interest between the clinical groups.

To systematically assess enriched gene sets between the clinical groups including the ‘complete remission’ status, all gene set collections were combined into one gene matrix transposed file (gmt) as input for GSEA, which was then performed as stated above. Top 100 enriched (NES) and significant (p < 0.05) scores were selected per corresponding cell type, translated into a ModuleScore and tested for significance between the clinical and CR states using paired Wilcoxon signed rank test.

#### GO overrepresentation analysis

To identify enriched terms among the DEGs from the MAST analyses, GO overrepresentation analysis was performed with the clusterProfiler R package (v3.16.1, [58]). The function `enricher()` was used to run GO analysis based on the same gmt files as used for GSEA.

#### Surfaceome filtering

DEGs from the MAST comparison of aberrant-like cells from patients at initial diagnosis against healthy-like cells from healthy donors within the memory CD8+ T cell subset were filtered for surface proteins using Cell Surface Protein Atlas data including validated surfaceome proteins [59]. Briefly, surface proteins annotated in Table A of the file http://wlab.ethz.ch/cspa/data/S2_File.xlsx (21 Apr 2021) were filtered for the category ’1 - high confidence’ and DEGs were filtered for the intersection with the remaining gene symbols in the surfaceome table.

#### Cell - cell interaction analysis

Cell-cell interactions were inferred with CellphoneDB2.0 [60] using normalized and decontX corrected count data of the human bone marrow data set. Receptor-ligand interactions were inferred for mean expression within each cell label cluster as well as for clusters having the combined information of cell label and DA-Seq information. For downstream analyses, significant interactions with an adjusted p-value < 0.05 were considered, which required an expression of receptor and ligand in at least 10% of the cells per cluster. CellphoneDB2.0 was computed per patient and the significant interaction counts were grouped over the respected disease subgroups.

#### RNA Velocity

To investigate developmental dynamics, scVelo (v0.2.4, [61]) in combination with Velocyto (v0.17.17, [62]) in Python (v3.9.7) was used. Reads were annotated as spliced, unspliced and ambiguous. The pipeline was run individually for each sample and data from resulting loom files were combined. Cells were subsetted based on prior analysis of CD8+ T cells. Splicing kinetics were recovered using `recover_dynamics()` with standard parameters, velocities were computed using velocity (mode=`’dynamical’)` and the velocity graph was calculated by `velocity_graph()` with standard parameters. Finally, for visualization, summarized velocity vectors are plotted using the `velocity_embedding_stream()` function in UMAP space in combination with the dissimilarity score. For plotting of single marker expression `velocity()` was used.

### Bulk RNA-sequencing and TCR clonotyping

#### BM preparation, staining and sorting

For sorting of CXCR3+ and CXCR3-cells, BM and PB samples of MM patients were thawed and processed as described above. Cells were stained with CD3-APCCy7, CD4-BUV737, CD8-BUV395 and CXCR3-PECy7 antibody. For sorting of CXCR3+ and CXCR3-cells, single, live CD3+CD4-CD8+ cells were gated and sorted as CXCR3-or CXCR3+ cells, respectively. 1000 CXCR3+ and CXCR3-CD8+ T cells from each sample were sorted on FACSAria Fusion equipped with a 100 µm nozzle.

#### Bulk RNA-sequencing and gene expression analysis

RNA was isolated using PicoPure RNA Isolation Kit (ThermoFisher), bulk RNA-sequencing libraries were generated using the SMART Seq Stranded Total RNA-Seq kit (Takara) and sequenced using the Illumina NovaSeq 6000 platform (2 x 100 bp). Adapter trimming was performed with Skewer (v0.2.2, [63]). Reads were aligned to human reference GRCh38 using STAR (v2.5.2b, [64]) and gene count tables were generated using Gencode v.32 annotations. Differential expression between samples was tested using the R/Bioconductor package DESeq2 (v1.30.1, [65]). Sample origin (BM vs. PB) was added to the design formula (condition: CXCR3+ vs. CXCR3-CD8+ T) to retrieve significantly upregulated genes for CXCR3+ CD8+ T cells within the BM (termed bulkRNA Remodeling Module).

#### TCR clonotype analysis

Analysis and quantification of the TCR receptor profiles, statistical analysis, and visualization were performed using three main tools: MiXCR (v3.0.13, [66]), VDJtools (v1.2.1, [67]) and immunarch (v0.6.6, [68]). Raw bulk RNA sequencing data of sorted CD8+ T cells in FASTQ format was used as the input for the TCR clonotype analysis. Analyze shotgun command of MiXCR was used to align variable (V), diversity (D), joining (J), and constant (C) genes of T-cell receptors, correct PCR and sequencing errors, assemble bulk RNA-seq reads by CDR3 region to the reference IMGT [69] library and export bulk TCR clonotypes. To this end, the default parameters recommended by the developers for RNA-seq data were used. Basic analysis, diversity estimation and repertoire overlap analysis modules of VDJtools were then used for the downstream analysis of the bulk TCR clonotypes provided by the MiXCR output. For the TCR repertoire clonality comparison between groups, the clonality metric was calculated as [1 – normalized Shannon Wiener Diversity Index] [70]. Significant differences were evaluated by paired Wilcoxon signed rank test. For TCR repertoire overlap quantification, the Jaccard index was utilized. Hierarchical clustering of the quantified TCR repertoire overlap was then performed using `hclust()` function of the R MASS package. `JoinSamples()` command of VDJtools were used to highlight overlapping TCR clonotypes by representative CDR3 amino acid sequence between CXCR3 status and sample origin (BM, PB) of CD8+ T cells of single patients. Additionally, the frequency of the top 10 most abundant TCR clonotypes across different samples was tracked using immunarch. Clonotype tracking was performed by the representative CDR3 amino acid sequence of TCR clonotypes.

### T cells in vitro cytokine assay

CD3+ T cells were enriched from the BM of 30 MM patients using the Pan T cell isolation kit with MS columns (both Miltenyi Biotec, Bergisch, Germany). 5 x 10^5^ CD3+ T cells were plated in 0.5 ml T cell expansion medium (Stemcell Technologies, Cologne, Germany) with 50 IE/ml IL-2, 1 % Pen/Strep (both from Sigma-Aldrich, Taufkirchen, German) in 24 well-plates and incubated overnight at 37 °C, 5 % CO2. On the next day, GolgiStop (0.66 µl/ml) (both BD Biosciences, Heidelberg, Germany) was added and cells were stimulated with PMA (50 ng/ml) and Ionomycin (1 µg/ml) (both Sigma-Aldrich, Taufkirchen, Germany). 6 h after incubation, intra-cellular staining was performed using transcription factor buffer set (BD Biosciences, Heidelberg, Germany) according to manufacturer’s instructions. Briefly, cells were washed twice in PBS and stained with cell surface antibodies for 20 min at 4 °C. Subsequently, antibody-conjugated cells were fixed and permeabilized for intracellular staining before washed twice with 1x Perm/Wash buffer and stained with antibodies against intra-cellular markers at 4 °C for 45 min. Cells were washed twice with 1x Perm/Wash buffer and measurements were acquired on cell analyzer FACS Lyrics (BD Bioscience, Heidelberg, Germany). Controls without PMA and Ionomycin stimulation were included in this assay. Flow cytometry data were visualized in FlowJo (Treestar).

### Multiplex Immunofluorescence

The frequency, localization and spatial proximity of T cell subpopulations and plasma cells, as well as their expression of respective markers LAT1 and CXCR3 was analyzed by multispectral imaging (MSI). Formalin fixed and paraffin embedded (FFPE) bone marrow (BM) biopsies of patients with MM (n=33), and control BM tissue of patients with B cell Non-Hodgkin lymphomas without evidence for BM infiltration (n=12) and myelodysplastic syndromes were collected between 2017 and 2020 at the Institute of Pathology of the Medical Faculty of the Martin-Luther University Halle-Wittenberg, Germany. The use of FFPE tissue samples was approved by the Ethical Committee of the Medical Faculty of the Martin Luther University Halle-Wittenberg, Halle, Germany (2017-81). The staining procedure was performed as recently described [71]. The marker panel used for staining included mAb directed against CD3 (Labvision. Germany. clone SP7), CD8 (Abcam, Cambridge, UK, clone SP16), MUM1 (Dako, USA, clone MUM1p), LAT1 (Abcam, Cambridge, UK, clone EPR17573) and CXCR3 (Abcam, Cambrdige, UK, clone ab133420).

Briefly, all primary mAb were incubated for 30 min. Tyramide signal amplification (TSA) visualization was performed using the Opal seven-color IHC kit containing fluorophores Opal 520, Opal 540, Opal 570, Opal 620, and Opal 690 (Perkin Elmer Inc., Waltham, MA, USA), and DAPI. Stained slides were imaged employing the PerkinElmer Vectra Polaris platform. Cell segmentation and phenotyping of the cell subpopulations were performed using the inForm software (PerkinElmer Inc., USA). The frequency of all immune cell populations analyzed and the cartographic coordinates of each stained cell type were obtained. The spatial distribution of cell populations was analyzed using an R script for immune cell enumeration and relationship analysis.

## Data availability statement

Single-cell RNA-seqquencing and bulk RNA-sequencing data are available at the European Genome-phenome Archive (EGA) under accession number EGAS00001006980.

**Table 2:**
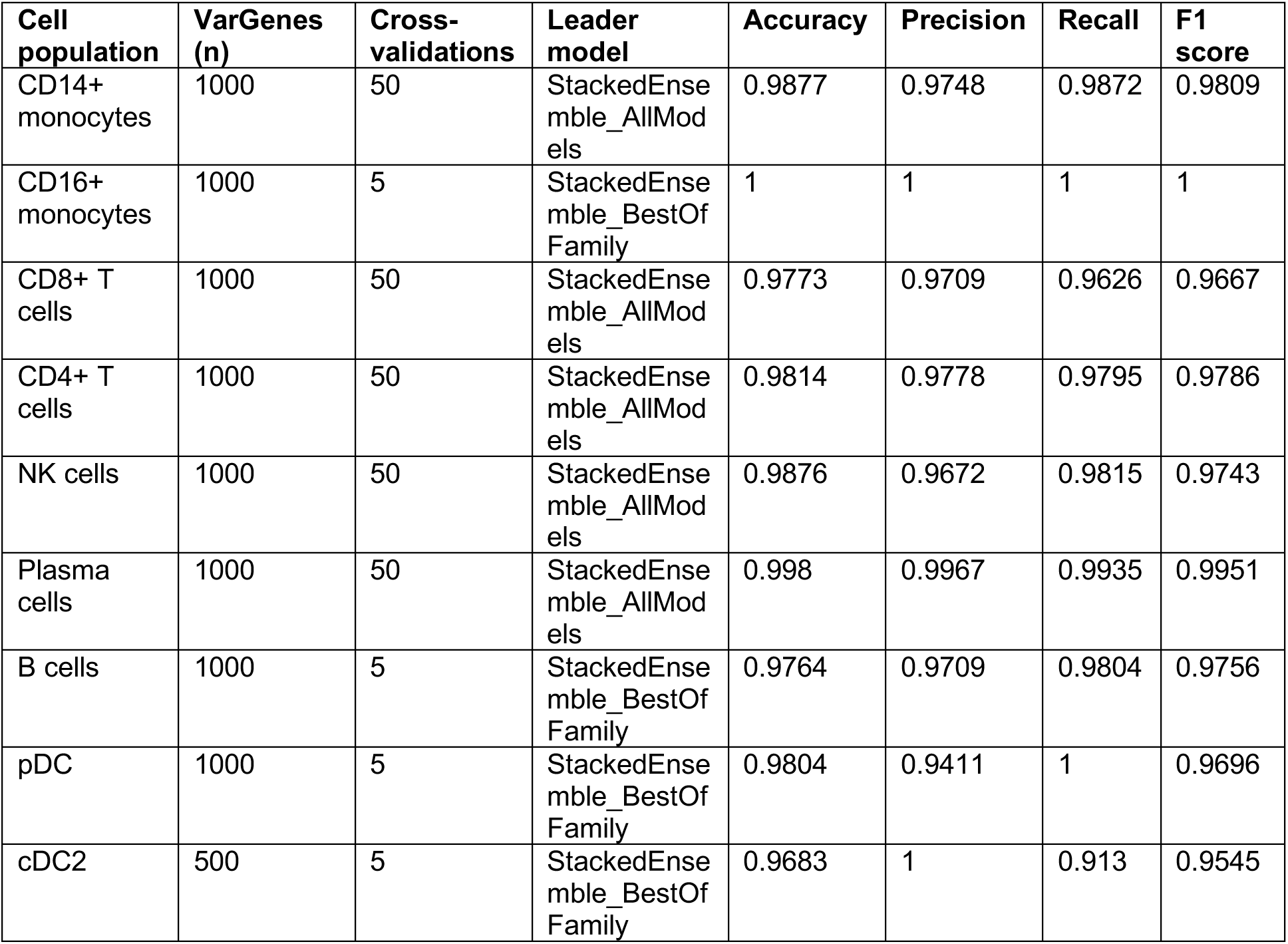
Input parameters, chosen model and prediction statistics for aberrant cell type classification using H2O (see methods)

## Notes

### Competing Interest Statement

The authors have declared no competing interest.

## References

1. Noy, R. and J.W. Pollard, Tumor-associated macrophages: from mechanisms to therapy. Immunity, 2014. 41(1): p. 49–61.

2. Thommen, D.S. and T.N. Schumacher, T Cell Dysfunction in Cancer. Cancer Cell, 2018. 33(4): p. 547–562.

3. van der Leun, A.M., D.S. Thommen, and T.N. Schumacher, CD8(+) T cell states in human cancer: insights from single-cell analysis. Nat Rev Cancer, 2020. 20(4): p. 218–232.

4. Veglia, F., M. Perego, and D. Gabrilovich, Myeloid-derived suppressor cells coming of age. Nat Immunol, 2018. 19(2): p. 108–119.

5. Veglia, F., E. Sanseviero, and D.I. Gabrilovich, Myeloid-derived suppressor cells in the era of increasing myeloid cell diversity. Nat Rev Immunol, 2021. 21(8): p. 485–498.

6. Fonseca, D.M., et al., Microbiota-Dependent Sequelae of Acute Infection Compromise Tissue-Specific Immunity. Cell, 2015. 163(2): p. 354–66.

7. Roquilly, A., et al., Alveolar macrophages are epigenetically altered after inflammation, leading to long-term lung immunoparalysis. Nat Immunol, 2020. 21(6): p. 636–648.

8. de Jong, M.M.E., et al., The multiple myeloma microenvironment is defined by an inflammatory stromal cell landscape. Nat Immunol, 2021. 22(6): p. 769–780.

9. Ghobrial, I.M., et al., The bone-marrow niche in MDS and MGUS: implications for AML and MM. Nat Rev Clin Oncol, 2018. 15(4): p. 219–233.

10. Dutta, A.K., et al., Single-cell profiling of tumour evolution in multiple myeloma - opportunities for precision medicine. Nat Rev Clin Oncol, 2022. 19(4): p. 223–236.

11. Ledergor, G., et al., Single cell dissection of plasma cell heterogeneity in symptomatic and asymptomatic myeloma. Nat Med, 2018. 24(12): p. 1867–1876.

12. Tirier, S.M., et al., Subclone-specific microenvironmental impact and drug response in refractory multiple myeloma revealed by single-cell transcriptomics. Nat Commun, 2021. 12(1): p. 6960.

13. Liu, R., et al., Co-evolution of tumor and immune cells during progression of multiple myeloma. Nat Commun, 2021. 12(1): p. 2559.

14. Zavidij, O., et al., Single-cell RNA sequencing reveals compromised immune microenvironment in precursor stages of multiple myeloma. Nat Cancer, 2020. 1(5): p. 493–506.

15. Das, R., et al., Microenvironment-dependent growth of preneoplastic and malignant plasma cells in humanized mice. Nat Med, 2016. 22(11): p. 1351–1357.

16. Lehners, N., et al., Analysis of long-term survival in multiple myeloma after first-line autologous stem cell transplantation: impact of clinical risk factors and sustained response. Cancer Med, 2018. 7(2): p. 307–316.

17. Paquin, A., et al., Characteristics of exceptional responders to autologous stem cell transplantation in multiple myeloma. Blood Cancer J, 2020. 10(8): p. 87.

18. Arteche-Lopez, A., et al., Multiple myeloma patients in long-term complete response after autologous stem cell transplantation express a particular immune signature with potential prognostic implication. Bone Marrow Transplant, 2017. 52(6): p. 832–838.

19. Bryant, C., et al., Long-term survival in multiple myeloma is associated with a distinct immunological profile, which includes proliferative cytotoxic T-cell clones and a favourable Treg/Th17 balance. Blood Cancer J, 2013. 3: p. e148.

20. Pessoa de Magalhaes, R.J., et al., Analysis of the immune system of multiple myeloma patients achieving long-term disease control by multidimensional flow cytometry. Haematologica, 2013. 98(1): p. 79–86.

21. Greipp, P.R., et al., International staging system for multiple myeloma. J Clin Oncol, 2005. 23(15): p. 3412–20.

22. Diaz-Tejedor, A., et al., Immune System Alterations in Multiple Myeloma: Molecular Mechanisms and Therapeutic Strategies to Reverse Immunosuppression. Cancers (Basel), 2021. 13(6).

23. Broyl, A., et al., Gene expression profiling for molecular classification of multiple myeloma in newly diagnosed patients. Blood, 2010. 116(14): p. 2543–53.

24. Swamydas, M., et al., Deciphering mechanisms of immune escape to inform immunotherapeutic strategies in multiple myeloma. J Hematol Oncol, 2022. 15(1): p. 17.

25. Susek, K.H., et al., The Role of CXC Chemokine Receptors 1-4 on Immune Cells in the Tumor Microenvironment. Front Immunol, 2018. 9: p. 2159.

26. Zheng, L., et al., Pan-cancer single-cell landscape of tumor-infiltrating T cells. Science, 2021. 374(6574): p. abe6474.

27. Andreatta, M., et al., Interpretation of T cell states from single-cell transcriptomics data using reference atlases. Nat Commun, 2021. 12(1): p. 2965.

28. Cibrian, D. and F. Sanchez-Madrid, CD69: from activation marker to metabolic gatekeeper. Eur J Immunol, 2017. 47(6): p. 946–953.

29. Te Boekhorst, P.A., et al., T-lymphocyte reconstitution following rigorously T-cell-depleted versus unmodified autologous stem cell transplants. Bone Marrow Transplant, 2006. 37(8): p. 763–72.

30. Schlenke, P., et al., Immune reconstitution and production of intracellular cytokines in T lymphocyte populations following autologous peripheral blood stem cell transplantation. Bone Marrow Transplant, 2001. 28(3): p. 251–7.

31. van der Velden, A.M., et al., Development of T cell-mediated immunity after autologous stem cell transplantation: prolonged impairment of antigen-stimulated production of gamma-interferon. Bone Marrow Transplant, 2007. 40(3): p. 261–6.

32. Hagiwara, K., et al., Dynamics of age-versus therapy-related clonal hematopoiesis in long-term survivors of pediatric cancer. Cancer Discovery, 2023.

33. Dangaj, D., et al., Cooperation between Constitutive and Inducible Chemokines Enables T Cell Engraftment and Immune Attack in Solid Tumors. Cancer Cell, 2019. 35(6): p. 885–900 e10.

34. Vilgelm, A.E. and A. Richmond, Chemokines Modulate Immune Surveillance in Tumorigenesis, Metastasis, and Response to Immunotherapy. Front Immunol, 2019. 10: p. 333.

35. Goodyear, O.C., et al., Neoplastic plasma cells generate an inflammatory environment within bone marrow and markedly alter the distribution of T cells between lymphoid compartments. Oncotarget, 2017. 8(18): p. 30383–30394.

36. Lutz, R., et al., Therapeutic Advances Propelled by Deciphering Tumor Biology and Immunology-Highlights of the 8th Heidelberg Myeloma Workshop. Cancers (Basel), 2021. 13(16).

37. Tokunaga, R., et al., CXCL9, CXCL10, CXCL11/CXCR3 axis for immune activation - A target for novel cancer therapy. Cancer Treat Rev, 2018. 63: p. 40–47.

38. Flores-Montero, J., et al., Next Generation Flow for highly sensitive and standardized detection of minimal residual disease in multiple myeloma. Leukemia, 2017. 31(10): p. 2094–2103.

39. Hao, Y., et al., Integrated analysis of multimodal single-cell data. Cell, 2021. 184(13): p. 3573–3587 e29.

40. Wang, Z., et al., Celda: a Bayesian model to perform co-clustering of genes into modules and cells into subpopulations using single-cell RNA-seq data. NAR Genom Bioinform, 2022. 4(3): p. lqac066.

41. Tickle, T.I., Georgescu, C., Brown, M. & Haas, B. inferCNV of the Trinity CTAT Project 2019; Available from:.

42. Plummer, M., JAGS: a program for analysis of Bayesian graphical models using gibbs sampling. 2003: Proc. 3rd International Workshop on Distributed Statistical Comptuting.

43. Gu, Z., R. Eils, and M. Schlesner, Complex heatmaps reveal patterns and correlations in multidimensional genomic data. Bioinformatics, 2016. 32(18): p. 2847–9.

44. Gu, Z., et al., circlize Implements and enhances circular visualization in R. Bioinformatics, 2014. 30(19): p. 2811–2.

45. Wickham, H., Seidel, D., scales: Scale Functions for Visualization. 2020.

46. Ooms, J. magick: Advanced Graphics and Image-Processing in R. R package version 2.7.3. 2021; Available from: https://CRAN.R-project.org/package=magick.

47. Team, T.I.D. ImageMagick. 2021; Available from:.

48. Aran, D., et al., Reference-based analysis of lung single-cell sequencing reveals a transitional profibrotic macrophage. Nat Immunol, 2019. 20(2): p. 163–172.

49. Zhao, J., et al., Detection of differentially abundant cell subpopulations in scRNA-seq data. Proc Natl Acad Sci U S A, 2021. 118(22).

50. LeDell, E., Poirier, S., H2o automl: Scalable automatic machine learning. Proceedings of the AutoML Workshop at ICML, 2020.

51. Kuhn, M., Building Predictive Models in R Using the caret Package. Journal of Statistical Software, 28(5), 1–26, 2008.

52. Regev, A., et al., The Human Cell Atlas. Elife, 2017. 6.

53. Hie, B., B. Bryson, and B. Berger, Efficient integration of heterogeneous single-cell transcriptomes using Scanorama. Nat Biotechnol, 2019. 37(6): p. 685–691.

54. Finak, G., et al., MAST: a flexible statistical framework for assessing transcriptional changes and characterizing heterogeneity in single-cell RNA sequencing data. Genome Biol, 2015. 16: p. 278.

55. Subramanian, A., et al., Gene set enrichment analysis: a knowledge-based approach for interpreting genome-wide expression profiles. Proc Natl Acad Sci U S A, 2005. 102(43): p. 15545–50.

56. Liberzon, A., et al., The Molecular Signatures Database (MSigDB) hallmark gene set collection. Cell Syst, 2015. 1(6): p. 417–425.

57. Korotkevich, G., Sukhov, V., Budin, N., Shpak, B., Artyomov M.N., Sergushichev, A., Fast gene set enrichment analysis. bioRxiv, 2021.

58. Yu, G., et al., clusterProfiler: an R package for comparing biological themes among gene clusters. OMICS, 2012. 16(5): p. 284–7.

59. Bausch-Fluck, D., et al., A mass spectrometric-derived cell surface protein atlas. PLoS One, 2015. 10(3): p. e0121314.

60. Efremova, M., et al., CellPhoneDB: inferring cell-cell communication from combined expression of multi-subunit ligand-receptor complexes. Nat Protoc, 2020. 15(4): p. 1484–1506.

61. Bergen, V., et al., Generalizing RNA velocity to transient cell states through dynamical modeling. Nat Biotechnol, 2020. 38(12): p. 1408–1414.

62. La Manno, G., et al., RNA velocity of single cells. Nature, 2018. 560(7719): p. 494–498.

63. Jiang, H., et al., Skewer: a fast and accurate adapter trimmer for next-generation sequencing paired-end reads. BMC Bioinformatics, 2014. 15: p. 182.

64. Dobin, A., et al., STAR: ultrafast universal RNA-seq aligner. Bioinformatics, 2013. 29(1): p. 15–21.

65. Love, M.I., W. Huber, and S. Anders, Moderated estimation of fold change and dispersion for RNA-seq data with DESeq2. Genome Biol, 2014. 15(12): p. 550.

66. Bolotin, D.A., et al., MiXCR: software for comprehensive adaptive immunity profiling. Nat Methods, 2015. 12(5): p. 380–1.

67. Shugay, M., et al., VDJtools: Unifying Post-analysis of T Cell Receptor Repertoires. PLoS Comput Biol, 2015. 11(11): p. e1004503.

68. ImmunoMindTeam, immunarch: an R package for painless bioinformatics analysis of T-cell and B-cell immune repertoires. 2019.

69. Lefranc, M.P., et al., IMGT unique numbering for immunoglobulin and T cell receptor variable domains and Ig superfamily V-like domains. Dev Comp Immunol, 2003. 27(1): p. 55–77.

70. Tumeh, P.C., et al., PD-1 blockade induces responses by inhibiting adaptive immune resistance. Nature, 2014. 515(7528): p. 568–71.

71. Bauer, M., et al., Multiplex immunohistochemistry as a novel tool for the topographic assessment of the bone marrow stem cell niche. Methods Enzymol, 2020. 635: p. 67–79.

