## Supplementary material for "Multiple myeloma long-term survivors display sustained immune alterations decades after first line therapy"

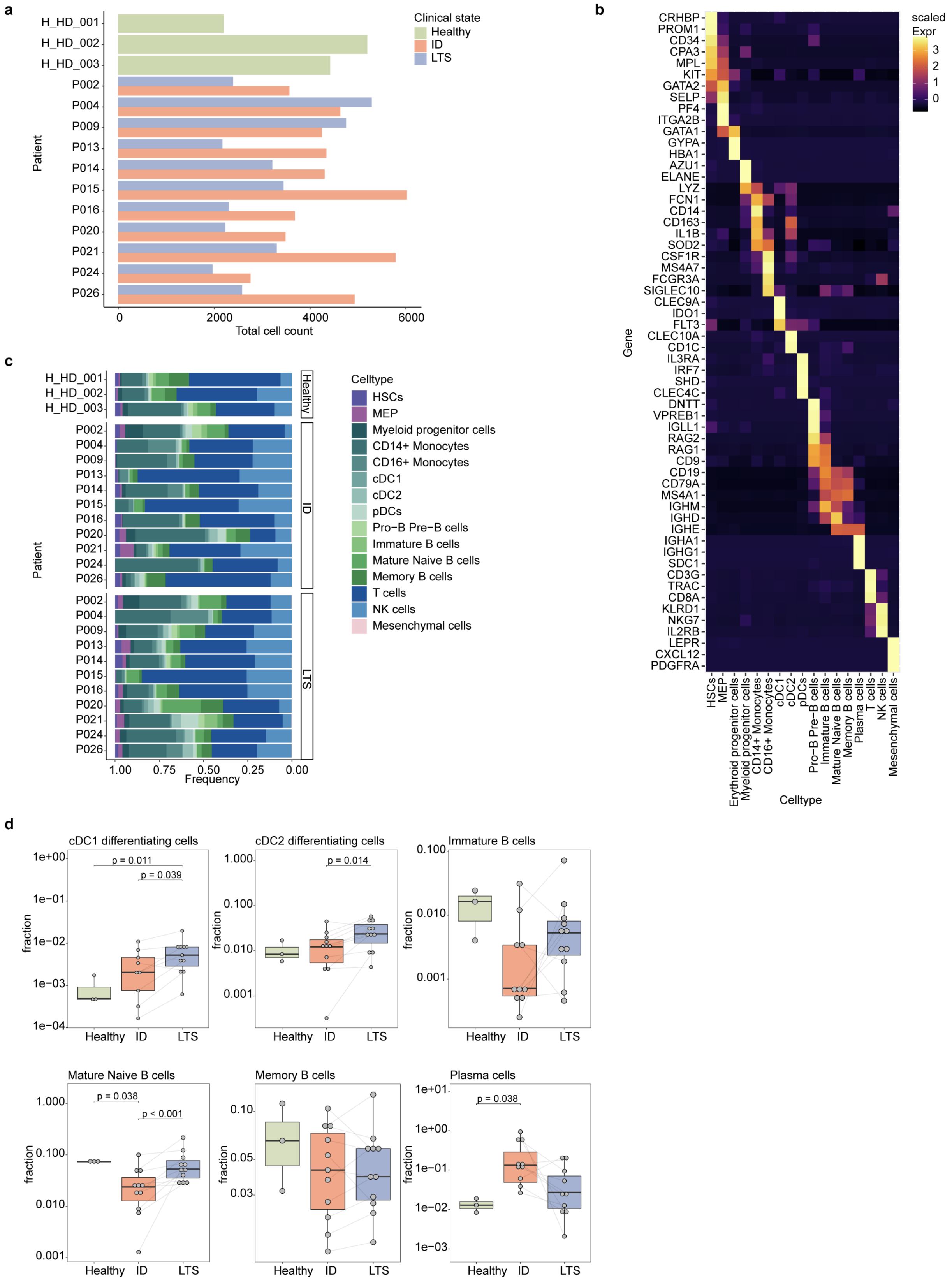


**Extended data Figure 1. Single-cell RNA-sequencing of the bone marrow ecosystem of multiple myeloma long-term survivor patients.**
**(a)** Total cell counts per sample and clinical state (Healthy; ID: initial diagnosis; LTS: long-term survival). **(b)** Gene expression heatmap of major marker genes for individual cell types; average gene expression per cell type, scaled row-wise for each gene. **(c)** BM cell type composition for healthy controls and MM patients per clinical condition (erythroid progenitors and plasma cells were excluded due to high

variation between sample and clinical state). **(d)** Differential proportion analysis (cell type fraction of total BM cells) for conventional dendritic cells 1 and 2 (cDCs), differentiated B cell compartment (immature, mature, memory, plasma cells). Significance was tested using Wilcoxon rank sum test for unpaired comparison between healthy (n=3) and ID (n=11), or by paired Wilcoxon signed rank test between ID and LTS (n=11).

Abbreviations: HSCs: hematopoietic stem cells, MEP: megakaryocyte-erythrocyte progenitors, MyeloP: myeloid progenitors, cDC1/2: conventional dendritic cells 1/2, pDCs: plasmacytoid dendritic cells, NK: natural killer cells, MSCs: mesenchymal stem cells; ID: initial diagnosis, LTS: long-term survival.

Box plots: center line, median; box limits, first and third quartile; whiskers, smallest/largest value no further than 1.5*IQR from corresponding hinge; dots: cell type fraction of total BM cells of each sample.

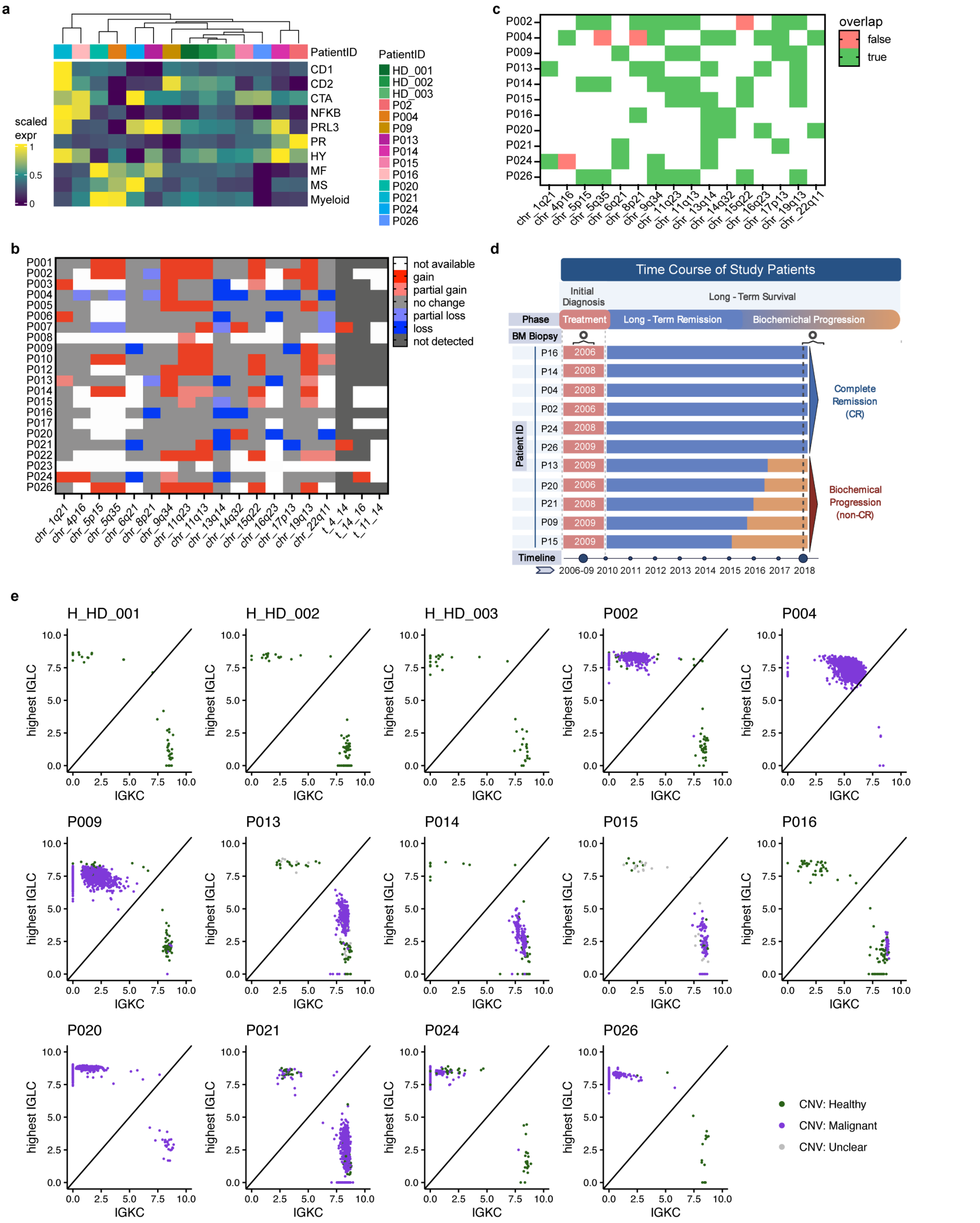


**Extended data Figure 2.1. Analyses of the plasma cell compartment during myeloma long-term survival.**
**(a)** Heatmap showing average expression patterns (module scores; scaled per score) of known bulk RNA-sequencing signatures per patient's plasma cells. Samples are ordered by Euclidean distance. **(b)** Copy number aberrations (CNAs) within plasma cells of each patient at ID detected by clinical routine FISH analysis. **(c)** Overlap of CNAs between standard FISH analysis and results from inferCNV (59 of 63 matches). **(d)** Time course of study patients subjected to scRNAseq from ID throughout LTS. **(e)** Scatterplots of immunoglobulin expression (highest lambda chain (IGLC) versus kappa chain (IGKC)) of healthy (green) and malignant (violet) plasma cells.

Abbreviations: ID: initial diagnosis; LTS: long-term survival; CAN: copy number aberrations; FISH: fluorescence in situ hybridization; IGLC: immunoglobulin light chain; LC: lambda chain; KC: kappa chain

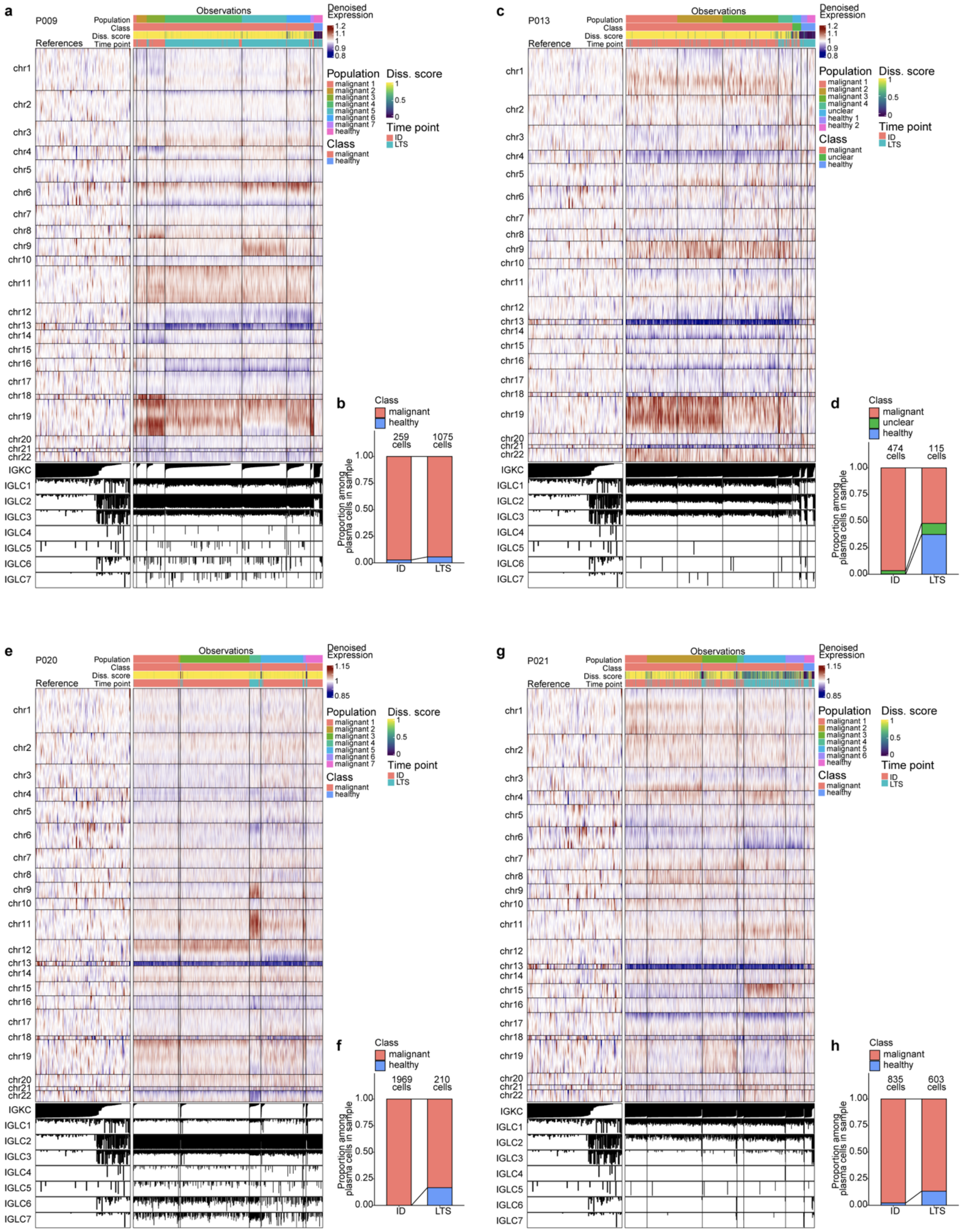


**Extended data Figure 2.2: Paired copy number aberration (CNA) analyses of the plasma cell compartment of MM LTS (P009, P013, P020, P021)**

**(a), (c), (e), (g)** inverCNV-based CNA heatmaps of denoised gene expression per within plasma cells of patients P009, P013, P020 and P021, respectively, compared to plasma cells from healthy controls (see methods). Only patients with sufficient numbers of PCs at ID and LTS states are indicated. Immunoglobulin light chain expression, dissimilarity score, subclonal annotation, malignancy class and clinical state are highlighted for each cell. **(b), (d), (f), (h)** Overall proportion of malignant (red) versus healthy (blue) plasma cells per patient (P009, P013, P020 and P021, respectively) as evaluated by inferCNV (methods).

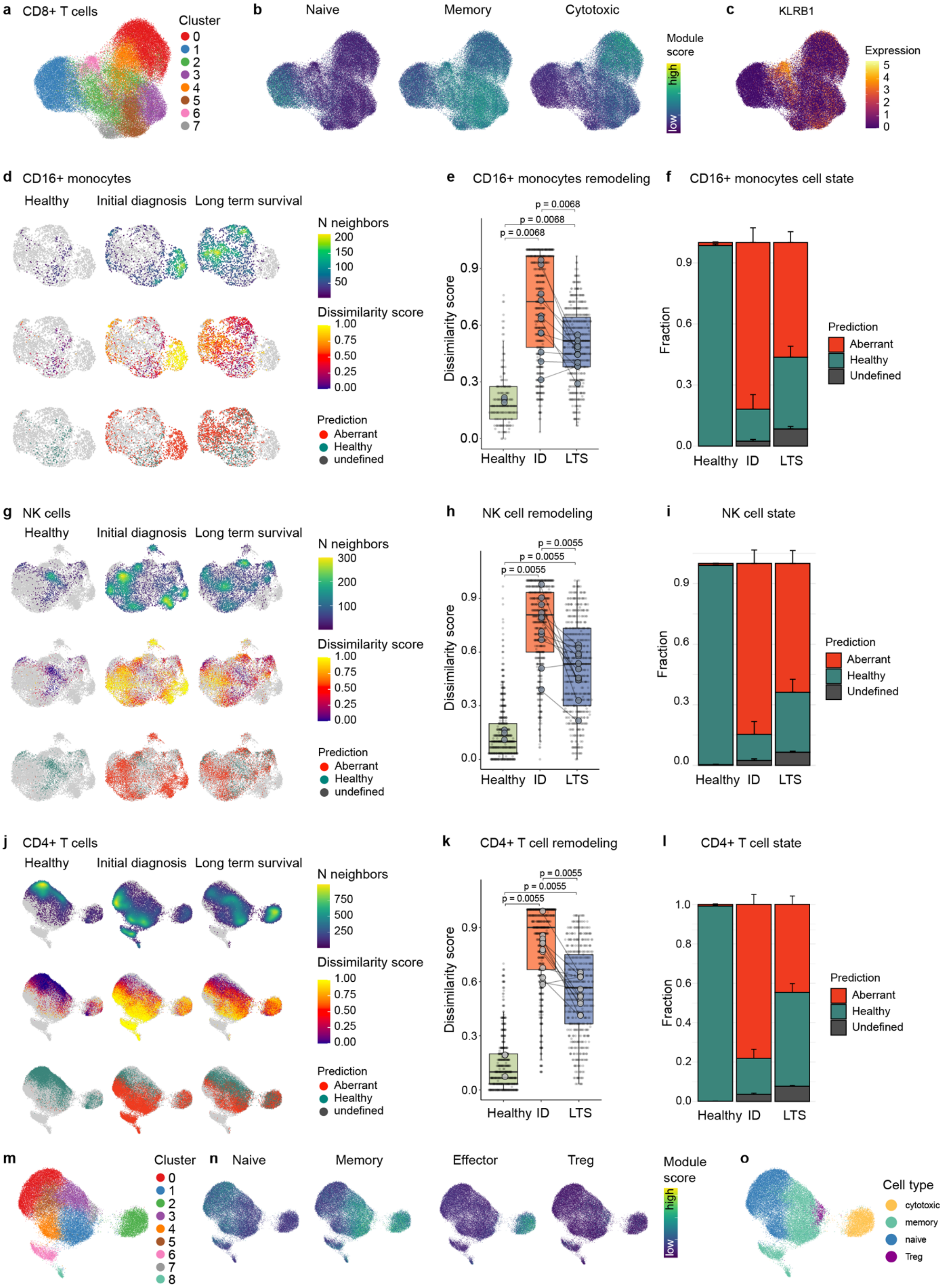


**Extended data Figure 3. Multiple myeloma long-term survivor patients display sustained signs of immune remodeling decades after a single therapy line.**
**(a-c)** CD8+ T cell dataset colored by (a) graph-based clusters, (b) module scores for naive, memory and cytotoxic CD8+ T cell gene signatures and (c) KLRB1 expression. **(d)** UMAP split by clinical groups showing cell density, dissimilarity scores and cell state predictions for CD16+ monocytes. Remaining cells from the corresponding other clinical groups are grayed out. **(e)** Boxplot of dissimilarity scores

summarized by clinical groups from d. **(f)** Fractions of predicted cell states by clinical group from d. **(g-l)** Similar visualizations as in (d,e,f) are shown for NK cells (g,h,i) and CD4+ T cells (j,k,l). **(m-o)** CD4+ T cell dataset colored by graph-based clusters (m), module scores for naive, memory and effector CD4+ T cell and Treg gene signatures (n) and CD4+ T cell subset classification based on clusters and module scores from (m,n) (o).

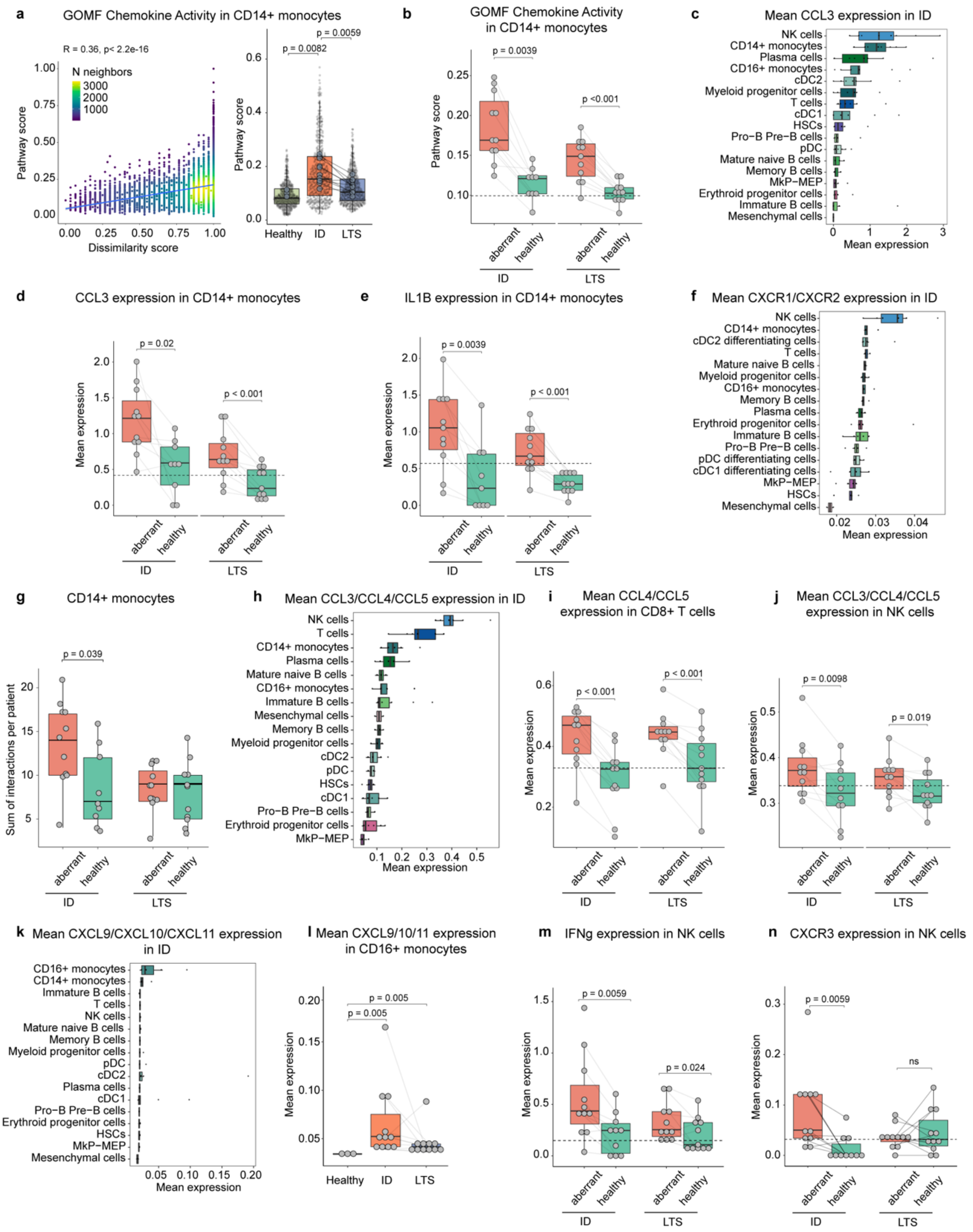


**Extended data Figure 4. An inflammatory circuit underlies immune remodeling during active disease and long-term survival.**

**(a)** Correlation of chemokine activity module score (GOMF chemokine activity) and dissimilarity score for CD14+ monocytes (left); boxplot of chemokine activity module score summarized by clinical groups (right). **(b)** Chemokine activity module score as in (a), but additionally split between predicted aberrant and healthy cells within the initial diagnosis and long-term survival groups. **(c)** CCL3 expression summarized by cell types at initial diagnosis. **(d,e)** Gene expression of CCL3 (d) and IL1B (e) in CD14+ monocytes split by clinical groups and cell state predictions. **(f)** Mean CXCR1/CXCR2 expression by cell types at initial diagnosis. **(g)** Number of predicted interactions of malignant plasma cells with CD14+ monocytes (interactome analysis, see methods) summarized by clinical group and cell state predictions. **(h)** Mean CCL3/CCL4/CCL5 expression by cell types at initial diagnosis. **(i)** Combined CCL4/CCL5 expression in CD8+ T cells grouped by clinical and cell state subsets. **(j)** Combined CCL3/CCL4/CCL5 expression in NK cells grouped by clinical and cell state subsets. **(k,l)** Mean CXCL9/CXCL10/CXCL11 expression plotted by cell types at initial diagnosis (k) and for clinical groups of the CD16+ monocyte subset (l). **(m,n)** Expression of IFNg (m) and CXCR3 (n) for NK cells split by clinical groups and predicted cell states.

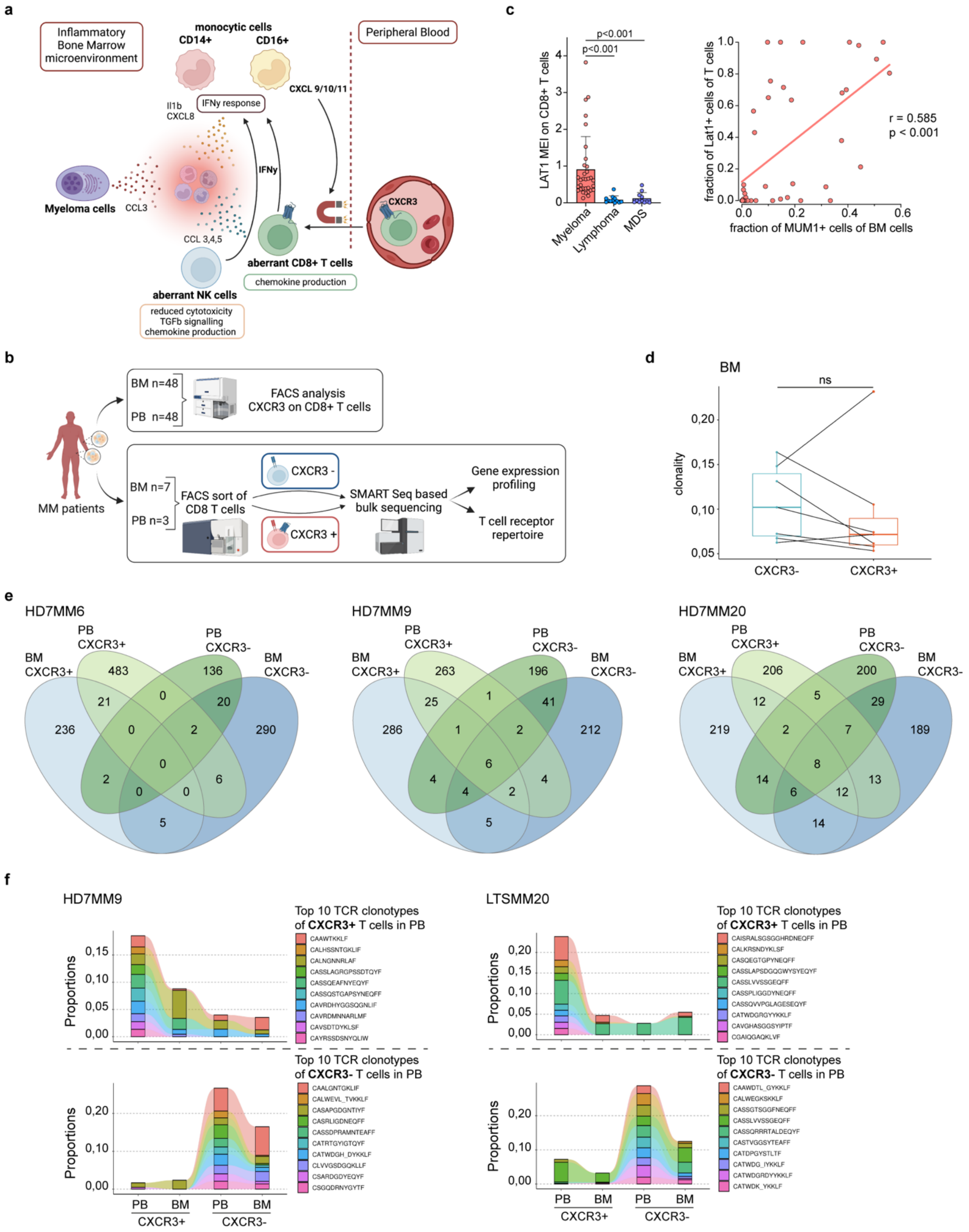


**Extended data Figure 5. Bone marrow infiltration of inflammatory T cells is associated with myeloma burden and serves as an accessible biomarker for disease activity.**
**(a)** Scheme illustrating the inflammatory circuit of aberrant immune cells in the BM microenvironment. **(b)** Study design scheme for the comparative analysis of CXCR3-positive and -negative CD8+ T cell subsets of MM patients in PB and BM to characterize myeloma associated T cells by flow cytometry (PB: n = 48, BM: n = 48) and their TCR repertoire and transcriptome by bulk RNAseq (PB: n = 3, BM: n = 7). **(c)** Left; LAT1 mean expression intensity (MEI) on BM CD8+ T cells during active disease state in MM, B cell non-Hodgkin lymphoma and MDS (as negative controls). Right; spearman correlation of LAT1 MEI with tumor burden measured by fraction of MUM1+ cells in the BM. Significance was tested by unpaired Wilcoxon rank sum test and corrected using BH for multiple comparison. **(d)** Boxplot

highlighting no significant differences in clonality of TCR repertoire between CXCR3- and CXCR3+ CD8+ T cells in the BM (n = 7 patients). Clonality metric was calculated as [1 – normalized Shannon Wiener Diversity Index]. Significant differences were evaluated by paired Wilcoxon signed rank test. **(e)** Venn diagram highlighting overlapping TCR clonotypes by representative CDR3 amino acid sequence between CXCR3 status and sample origin (BM, PB) of CD8+ T cells of individual patients. **(f)** Clonotype tracking by representative CDR3 amino acid sequence of shared clonotypes between the top 10 most abundant TCR clonotypes from CXCR3+ (top row) and CXCR3- (bottom row) peripheral blood (PB) CD8+ T cells across CXCR3+ or CXCR3- CD8+ T cell subsets in PB and BM.

Abbreviations: BM: bone marrow; PB: peripheral blood; MEI: mean expression intensity; MDS: myelodysplastic syndrome; ASCT: autologous stem cell transplantation; TCR: T cell receptor; BH: Benjamini-Hochberg

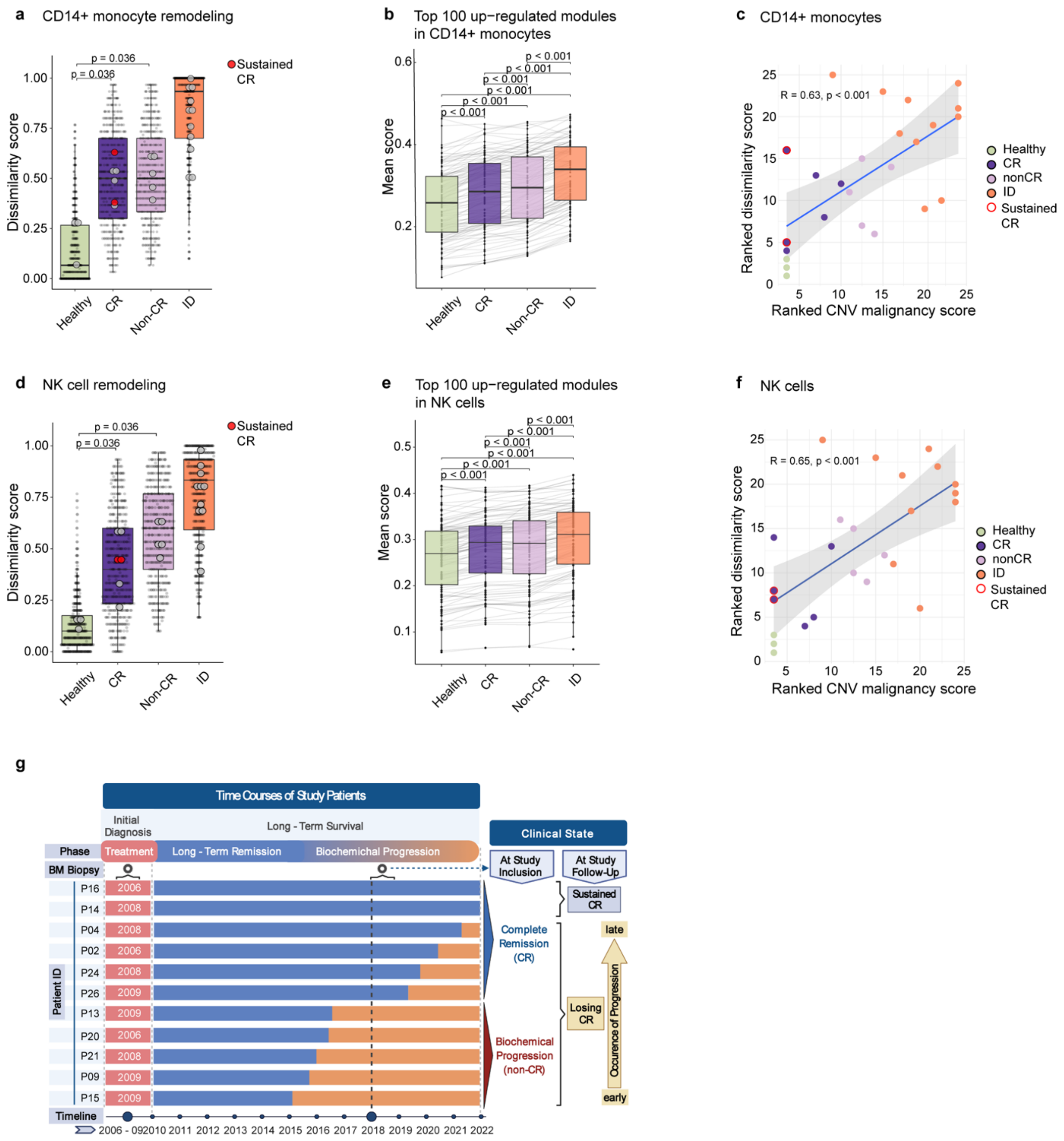


**Extended data Figure 6. Immune remodeling in LTS patients is associated with future disease resurgence and defective immune function even in the absence of measurable disease.** (**a**,**d**) Distribution of the dissimilarity score by clinical group with LTS patients split into CR (complete remission) and Non-CR, summarizing the remodeling of CD14+ monocytes **(a)** and NK cells **(d)**. Large dots indicate sample means. Sustained CR patients are highlighted in red. P-values from unpaired two-sided Wilcoxon rank-sum tests are shown. (**b**,**e**) Module Scores of top 100 upregulated pathways (ID vs. Healthy, see GSEA in methods) between clinical groups and CR status in CD14+ Monocytes (b) and NK cells (e). BH-corrected p values from paired one-tailed Wilcoxon signed rank test are highlighted. (**c**,**f**) Scatterplot of ranked mean dissimilarity score against CNV-based malignancy scores of CD14+ monocytes (c) and NK cells (f) for each patient and healthy controls; Spearman correlation was used to evaluate the relationship. **(g)** Clinical Follow up over 4 years: Time course of study patients subjected to scRNAseq from ID throughout LTS including study follow up to evaluate sustained CR.
Abbreviations: ID: initial diagnosis; LTS: long-term survival; CR: complete remission; NK: natural killer; BH: Benjamini Hochberg.
